# Mutations adjacent to the nucleotide-binding cleft of *Arabidopsis thaliana* ACTIN7 confer resistance to the actin-disrupting compound latrunculin B

**DOI:** 10.1101/2025.01.24.634782

**Authors:** J. Alan Sparks, Liang Sun, Sabrina Chin, Nolan Ramanjulu, Jiangqi Wen, Simon Gilroy, Elison B. Blancaflor, Bibi Rafeiza Khan

## Abstract

A forward genetic screen identified a dominant-negative *Arabidopsis thaliana* mutant resistant to growth inhibition caused by the actin-disrupting compound latrunculin B (LatB). Map-based cloning combined with whole-genome sequencing revealed that the mutant referred to here as *lbr1* for *<u>L</u>at<u>B r</u>esistant1* had a point mutation in the *AT5G09810* gene, which encodes the vegetative actin (ACT) isoform ACT7. The cytosine to thymine mutation in the second exon of *ACT7* of *lbr1* led to substitution of proline to serine at position 34 (P34S) adjacent to the nucleotide-binding cleft of the ACT7 protein. Confirmation that *ACT7* is the causal gene for the *lbr1* phenotype was achieved through transgenic complementation with *ACT7 wild type* (*ACT7_WT_*) and *ACT7 P34S* (*ACT7_P34S_*) constructs. *ACT7_P34_* also rescued the seedling developmental defects and conferred partial resistance to LatB in the recessive *act7-5* mutant. Furthermore, expressing a P34S mutation in ACT2 (ACT2_P34S_), another vegetative ACT isoform, conferred partial LatB resistance to wild type. Finally, site-directed mutagenesis of ACT7 amino acid residues forming putative hydrogen bonds with LatB, based on yeast and mammalian actin docking and structural analyses, reveals domains adjacent to the actin nucleotide-binding cleft crucial for LatB’s effects on the plant actin cytoskeleton.

**Highlight:** Characterization of the dominant-negative *lbr1* mutant uncovers amino acid residues in the actin protein crucial for latrunculin’s mechanism of action in plants.

## Introduction

The primary root, also known as the taproot, serves as the framework on which the entire root system is constructed. As the primary root elongates, it differentiates into various specialized tissues that support water and nutrient uptake and the synthesis of important growth-regulating hormones (Gifford *et al*., 2024). Root hairs, for example, are specialized cells that emerge from epidermal cells of primary and lateral roots. They significantly increase the root’s surface area, thus aiding in the absorption of water and essential minerals and promoting the plant’s overall nutrient uptake. Given the critical importance of the root system, understanding the complex molecular and genetic networks that regulate its growth and development will aid in designing strategies to optimize crop productivity and improve plant health (Tsang *et al*., 2024).

The actin cytoskeleton plays a significant role in root growth and development. It does so by modulating numerous cellular processes, including vesicular and organelle trafficking, secretion, endocytosis, and exocytosis, hormone transport, and root hair tip growth (Paez-Garcia *et al*., 2018). The actin cytoskeleton also participates in signaling events triggered by various internal and external stimuli (Mao *et al*., 2016; Zhu *et al*., 2016; Maeda *et al*., 2020; Henty-Ridilla *et al*., 2013; Leontovyčová *et al*., 2019). Plant genomes have multiple actin isoforms, with the *Arabidopsis thaliana* genome encoding eight *ACTIN (ACT)* genes (McLean *et al*., 1990; Meagher, 1991). Based on their sequence phylogeny and expression patterns, they are grouped into two major classes, namely, vegetative and reproductive (McDowell *et al*., 1996a; Meagher *et al*., 1999; Kandasamy *et al*., 2010). The vegetative class consists of *ACT2*, *ACT7,* and *ACT8*, while *ACT1*, *ACT3*, *ACT4*, *ACT11,* and *ACT12* are grouped into the reproductive class (McDowell *et al*., 1996a; Meagher *et al*., 1999; Kandasamy *et al*., 2010). Several studies show that the vegetative *ACTINS* are highly expressed in younger tissues, including inflorescences, sepals, petals, shoots, roots, seeds, and microspores, whereas reproductive *ACTINS* are preferentially expressed in mature pollen and ovules (An *et al*.,1996; McDowell *et al*.,1996b; Meagher *et al*., 1999; Gilliland *et al*., 2002). Because the three vegetative *ACTINS* are differentially expressed and play distinct roles in plant morphogenesis (Gilliland *et al*., 2003; Kandasamy *et al*., 2009), they are further subdivided into two subclasses. Subclass 1, consisting of *ACT2* and *ACT8,* is preferentially expressed in older tissues and regulates root hair growth and leaf morphology (Ringli *et al*., 2002; Nishimura *et al*., 2003; Kandasamy *et al*., 2009; Vaškebová *et al*., 2018). Subclass 2 *ACT7* expression is significantly influenced by hormones, such as auxin and ethylene, and other environmental stimuli. It is strongly expressed in young, developing tissues and organs, including the root meristem (An *et al*., 1996; Kandasamy *et al*., 2001; McDowell *et al*.,1996b; Meagher *et al*., 2000; Numata *et al*., 2022).

Studies of vegetative *A. thaliana ACT* mutants have been instrumental in deciphering the function of ACT isoforms in root development. For instance, transfer (T)-DNA insertion *ACT2* null mutants (e.g., *act2-1*, *act2-3,* and *act2-4*) and *ACT2* missense mutants (e.g., *der1-1*, *der1-2*, and *der1-3)* all showed defective root hair elongation phenotypes, highlighting the importance of ACT2 in tip growth (Ringli *et al*., 2002; Gilliland *et al*., 2002; Nishimura *et al*., 2003). Furthermore, the T-DNA insertion *ACT7* null mutants, *act7-1*, *act7-4*, and *act7-5*, show significant primary root elongation defects, with the *act7-5* mutant displaying the most dramatic reduction in root growth (Gilliland *et al*., 2003; Kandasamy *et al*., 2010). Additional phenotypic defects of *act7* mutants include delayed and reduced germination and altered cellular polarity in primary roots (Gilliland *et al*., 2003; Kandasamy *et al*., 2010). Examination of *act7* roots showed an increase in cell density in the transition zone and a decrease in the size of the proximal meristematic zone (Takatsuka *et al*., 2018).

In addition to mutant analysis, pharmacological compounds have been instrumental in elucidating the function of the actin cytoskeleton during plant development (Hou *et al*., 2003; Hou *et al*., 2004; Staiger *et al*., 2009; Rosero *et al*., 2013; Pospich *et al*., 2020). One such compound used extensively in plant biological studies is latrunculin B (Lat B), which is a marine toxin isolated from the Red Sea sponge *Negombata magnifica* (formerly *Latrunculin magnifica*; Morton *et al*., 2000). LatB sequesters actin monomers or binds to the ends of filamentous-actin (F-actin), disrupting its organization and inhibiting polymerization of globular (G)-actin monomers into F-actin (Morton *et al*., 2000). Nanomolar concentrations of LatB lead to root developmental defects that mirror those of vegetative *ACT* mutants, including the inhibition of primary root and root hair tip growth (Bibikova *et al*., 1999; Baluška *et al*., 2001a; Yoo *et al*., 2012).

The distinct seedling phenotypes caused by LatB prompted us to use it in forward genetic screens to identify *A. thaliana* mutants that are hypersensitive or resistant to its growth inhibitory effects. Characterization of the isolated mutants and cloning of their corresponding genetic lesions led to the identification of proteins with previously unreported functions in root development, actin organization, and endomembrane dynamics. For example, a recessive mutant that was hypersensitive to LatB (*hlb1*) was shown to be disrupted in a gene encoding a plant-specific tetratricopeptide repeat domain-containing protein that mediates exocytotic and endocytic protein trafficking during primary root and root hair tip growth (Sparks *et al*., 2016). Another LatB hypersensitive mutant originally called *hlb2* was disrupted in a gene encoding the beige and Chediak Higashi (BEACH) domain-containing protein, SPIRRIG (SPI). SPI accumulates in the tips of elongating root hairs, where it participates in maintaining tip-focused F-actin and modulates the stability of BRICK1, a component of the actin-related protein (Arp2/3) actin nucleating complex, during root hair initiation (Chin *et al*., 2021). Furthermore, *hlb3* was disrupted in a gene encoding a class II formin, which is a protein involved in maintaining F-actin bundling (Sun *et al*., 2019). Here, we characterize another mutant identified from a LatB forward genetic screen. This mutant is strongly resistant to the growth inhibitory effects of LatB (hereafter referred to as *lbr1* for *<u>L</u>at<u>B</u> Resistant1*). We found that *lbr1* contains a point mutation in the *ACT7* gene, leading to an antimorphic, dominant-negative *act7* mutant. Using *lbr1*, we identify critical amino acid residues adjacent to the vegetative ACT nucleotide-binding cleft that specify the mechanism of action of LatB on the plant actin cytoskeleton.

## Materials and methods

### Forward genetic screening on latrunculin B

The *lbr1* mutant was isolated through a forward genetic screen of ethyl methanesulfonate (EMS)-mutagenized M2 *A. thaliana* Columbia-0 (Col-0) seedlings (Lehle Seeds, Round Rock, TX) grown on LatB. Briefly, mutagenized seeds were surface sterilized in 70% ethanol for 5 min, followed by 10 min in 30% (v/v) sodium hypochlorite and rinsed four times with sterile deionized water. The medium for screening the EMS-mutagenized seeds consisted of 0.5 x Murashige and Skoog (MS) basal salt medium with vitamins (PhytoTech Labs, USA) and 1% (w/v) sucrose. The pH of the medium was adjusted to 5.7 with potassium hydroxide. Solid medium for planting was prepared by adding 1% agar (w/v) (Sigma-Aldrich, St. Louis MO, USA) and was autoclaved for 20 minutes. A stock solution of 10 mM LatB (Cayman Chemicals, Ann Arbor, MI, USA) in dimethyl sulfoxide (DMSO) was added to the medium after it had cooled to ∼60°C to obtain a final working LatB concentration of 100 nM and was poured into 10 x 10 cm square Petri dishes (Avantor/VWR, Radnor, PA, USA) to a thickness of ∼2.5 mm. Once the plates solidified, EMS-sterilized seeds were carefully plated in rows on MS medium plates (without LatB) using a fine-tip transfer pipette. The plates were stratified at 4°C for 48 hours, then transferred to a Conviron growth chamber and grown vertically for three days at 120 μmol m^−2^ s^−1^ light intensity supplied by fluorescent bulbs and set to a 14-h-light and 10-h-dark cycle. Temperature in the chamber was maintained at 23°C. Subsequently, all seedlings of uniform size were transferred to plates containing 100 nM LatB medium using sterile forceps, and the position of the tips of the primary roots were marked on the bottom face of the plates using a Sharpie. The plates were returned to the Conviron growth chamber, and the seedlings were allowed to grow for an additional five days. Seedlings with long primary roots relative to other seedlings were selected, transplanted to pots with Metromix 350, and grown in a greenhouse at 23°C, with a 16-hour photoperiod of 150 μE m-2 sec-1 intensity until seeds were obtained. The phenotypes of progeny were confirmed by rescreening the seedlings for primary roots that were longer than those of wild type when grown in Petri dishes with 100 nM LatB.

### Map-based cloning and whole genome sequencing

To identify the *LBR1* gene, *lbr1* plants were backcrossed and those determined to be homozygous were outcrossed to the Landsberg erecta ecotype. To select seedlings for map-based cloning, outcrossed F_2_ seeds were sterilized, plated on MS-only medium, and grown in the same Conviron growth chamber used for screening. Three-day-old seedlings were transferred to plates with 100 nM LatB medium and grown for an additional 5 days. Seedlings with short primary roots similar to the wild type on the 100 nM LatB medium were selected for mapping. Sequence length polymorphism and cleaved amplified polymorphic sequence markers were used to map the *lbr1* mutation to chromosome 5 within coordinates 2,142,759 bp – 3,122,929 bp. DNA was then extracted from homozygous *lbr1* seedlings, and whole genome sequencing was performed. Approximately 103 million 150 bp Hi-Seq illumina paired-end reads from *lbr1* were obtained via next-generation sequencing (NGS) technologies. Whole genome NGS data from *hlb3* were used as a control sample (Sun *et al*., 2019). All NGS data were trimmed via Trimmomatic 0.35 (LEADING:3 TRAILING:3 SLIDINGWINDOW:4:20 MINLEN:36). The high-quality data after trimming were aligned to the *Arabidopsis thaliana* reference genome (TAIR10) via BWA MEM algorithm (Li and Durbin, 2009). All single nucleotide polymorphisms (SNPs) and insertion-deletions (Indels) were called using VarScan (v2.3.9) (Koboldt *et al*., 2012). To identify the unique SNPs and Indels for *lbr1*, a customized Python script was written to filter out homozygous SNPs and Indels that commonly occurred in the *hlb3* mutant. The unique SNPs and Indels in *lbr1* were annotated via ANNOVAR (Wang *et al*., 2010; Yang and Wang, 2015).

### Chemical treatments and seedling growth assays

For seedling growth assays, seeds of all genotypes were surface sterilized for 10 min in 30% (v/v) sodium hypochlorite and rinsed four times with sterile deionized water. Sterilized seeds were planted on 0.5 x MS basal salt medium with vitamins, 1% (w/v) sucrose, and 1% agar (w/v) and stratified at 4°C for 48 h. LatB was diluted in DMSO to a stock concentration of 10 mM, which was then used to prepare 0.5 x MS medium with various LatB concentrations. For primary root assays, 3-day-old seedlings that exhibited uniform growth were selected and transplanted onto 10 cm X 10 cm square Petri dishes containing either 0.5 x MS medium supplemented with an equivalent amount of DMSO (solvent controls) or with various concentrations of LatB. The position of the root tip upon transplant was marked with a black Sharpie on the back face of the Petri dish and seedlings were grown vertically for an additional six days in the same Conviron growth chamber under the same conditions used for forward genetic screening. For quantification of hypocotyl length, seeds were grown either on solvent control medium or LatB-supplemented medium, incubated in light for 12 hours, and then transferred to the dark and kept vertically for five days. Images of seedlings were captured using a Nikon Insight digital camera mounted on a copy stand.

To evaluate root hair length, wild type and *lbr1* seeds were planted on 48 × 64 mm coverslips coated with approximately a 1 mm thick layer of 0.5 x MS containing 1% (w/v) sucrose and 0.4% (w/v) Gelzan (Chai *et al*., 2022). The growth medium was supplemented with different concentrations of LatB, or with equivalent amounts of DMSO, as solvent controls, prior to layering onto coverslips. Coverslips were placed in 9 cm round Petri dishes, and seeds were grown for 4 to 5 days in a Conviron growth chamber. Root hairs were photographed with a Nikon SMZ1500 stereomicroscope (Nikon Instruments Inc., Melville, NY, USA). Primary root, hypocotyl, and root hair lengths were measured using the ImageJ v1.49 software (http://rsb.info.nih.gov/ij/).

### Quantification of F-actin density in the primary root elongation zone

To quantify F-actin in primary roots, wild type and *lbr1* seedlings expressing *UBQ10-GFP-ABD2-GFP* (Dyachok *et al*., 2014) were grown for three days on 10 x 10 cm square Petri dishes containing 0.5 x MS LatB-free medium, prior to transferring to MS medium with 100 nM LatB or solvent controls. After six days, seedlings were mounted on 48 × 64 mm coverslips and secured using a slab of 1% (w/v) Gelzan (Chai *et al*., 2022). Coverslips with seedlings were placed horizontally on the stage of an inverted Leica SP8-X point scanning confocal microscope (Leica Microsystems, Buffalo Grove, IL, USA) and imaged with a 63× water (numerical aperture = 1.20) immersion objective. Images of F-actin labeled with GFP in the primary root elongation zone were captured by illuminating seedlings with the 488 nm line of the SP8-X Argon laser and emission detected at 510 nm. F-actin density in wild type and *lbr1* roots was analyzed using the open source RhizoVision software originally developed for the quantification of root system architecture (Seethepalli *et al*., 2021). Grayscale confocal images of root cell F-actin in TIFF format were inverted using Photoshop (Adobe) and segmented using RhizoVision Explorer v2.0.3 (Seethepalli and York, 2020) and the pixel area occupied by filaments was divided by the total image pixel area (Supplementary Fig. S1).

### Generation of *ACT7* complementation constructs

The *proACT7:ACT7_WT_*, *proACT2:ACT2_WT_,* and *proACT7:ACT7_LBR1_* constructs were generated from full-length genomic (g) DNA. gDNA was extracted from 2-week-old wild type and *lbr1* mutant seedlings using the Plant DNAzol Reagent (ThermoFisher Scientific, Waltham, MA, USA). The full-length *ACT7_WT_* and *ACT7_LBR1_* gDNA, including the ∼1609 bp immediately upstream of the start codon (3021 bp total), were amplified using polymerase chain reaction (PCR) with the primers ACT7pro-F-HindIII (5′-CATAAGCTTGTCTTTTAGTGTGCATTCCTCAAA-3′) and ACT7pro-R-BamHI (5′-GTAGGATCCTTAGAAGCATTTCCTGTGAACAAT-3′). The full-length *ACT2_WT_* gDNA, including the ∼1634 bp immediately upstream of the start codon (2932 bp total), was PCR amplified using the primers ACT2pro-F-HindIII (5′-CATAAGCTTCACTTACATAGCGCATCGCAC-3′) and ACT2pro-R-BamHI (5′-GTAGGATCCTTAGAAACATTTTCTGTGAACGATTC-3′). The PCR products were cloned into a modified *pCAMBIA1390* vector (Wang *et al*., 2008).

### Site-directed mutagenesis of *ACT2* and *ACT7* and plant transformation

All the site-directed mutants were generated by primer extension with mutagenic primers in independent, nested PCRs, before combining them into the final product as previously described (Reikofski and Tao, 1992). The *ACT2_P34_*_S_, *ACT7_Y71G_*, *ACT7_D159G_, ACT7_T188G_, ACT7_E209G_,* and *ACT7_R211G_* mutants were generated from gDNA, extracted from 2-week-old wild type seedlings using the Plant DNAzol Reagent (ThermoFisher).

To mutate the cytosine located at position 100 from the start codon of the *ACT2* gDNA to a thymine, causing the codon to code for a serine instead of a proline, the SNP mutation was incorporated into the following primers: C-F (5′-GGCTGTTTTTTCCAGTGTTGT-3′) and B-R (5′-ACAACACTGGAAAAAACAGCC-3′). Two partial *ACT2* gDNA mutant products were amplified with the primer pairs A-F (5’-CTACCAGAATTTGGCTTGACCT-3’) and primer B-R; primers C-F and D-R (5’-TCAGAGCTCAGTTCAAATAATG-3’). Primers A-F and B-R yielded a partial *ACT2* gDNA product of 398 bp, while primers C-F and D-R yielded a partial *ACT2* gDNA product of 425 bp. Both mutated products were then used as templates for amplification with primers A-F and D-R, resulting in a partial *ACT2* mutant gDNA product of ∼802 bp. At the beginning of the partial ∼802 bp *ACT2* gDNA region, where primer A-F starts, there is a PflMI restriction site, while a BanII restriction site is at the end, where primer D-R ends. Therefore, the *pCAMBIA1390* vector with the *pACT2:ACT2_WT_* construct was digested with PflMI and BanII to remove the ∼802 bp from the *ACT2_WT_* construct, and this region was replaced with the mutant *ACT2* gDNA product of ∼802 bp to generate the *pACT2:ACT2_P34S_* construct.

Each of the *ACT7* point mutants was generated using the following primer pairs: Y71G-F (5’-TGAAGGGGCCAATCGAACAT-3’) and Y71G-R (5’-ATGTTCGATTGGCCCCTTCA-3’); D159G-F (5’-ATTCTGGTGGGGGTGTGTCT-3’) and D159G-R (5’-AGACACACCCCCACCAGAAT-3’); T188G-F (5’-TCGGGATCTCGGGGACTCAC-3’) and T188G-R (5’-GTGAGTCCCCGAGATCCCGA-3’); E209G-F (5’-TACCGCAGAACGGGGGATTGT-3’) and E209G-R (5’-ACAATCCCCCGTTCTGCGGTA-3’); R211G-F (5’-AAATTGTCGGGGACATAAAG-3’) and R211G-R (5’-CTTTATGTCCCCGACAATTT-3’).

Two partial *ACT7* gDNA mutant products were amplified to generate each of the *ACT7* point mutants. Thus, the *ACT7* gDNA was amplified with each of the mutant forward primers and primer V-R (5’-GGTGCTGAGGGATGCAAGGATTGATC-3’) and each of the reverse mutant primers and primer U-F (5’-CGTCCTAGGCACACTGGTGTCATGG-3’). For each of the *ACT7* point mutants, both mutated products were used as templates for amplification with primers U-F and V-R, yielding partial *ACT7* mutant gDNA products of 1032 bp. At the beginning of the partial 1032 bp *ACT7* gDNA region, where primer U-F starts, there is an AvrII restriction site, while a BbvCI restriction site is at the end, where primer V-R ends. Therefore, the *pCAMBIA1390* vector with the *pACT7:ACT7_WT_* construct was digested with AvrII and BbvCI to remove the 1032 bp from the *ACT7_WT_* construct, and this region was replaced with each of the mutant *ACT7* gDNA products of 1032 bp to generate the *pACT7:ACT7_Y71G_*, *pACT7:ACT7_D159G_, pACT7:ACT7_T188G_, pACT7:ACT7_E209G_,* and *pACT7:ACT7_R211G_* constructs. Sequencing to confirm mutagenesis was performed using DNASTAR Lasergene, version 12.1.

All transgenic lines were generated by the transformation of the constructs into *Agrobacterium tumefaciens* strain C58C1, followed by floral dip (Clough and Bent, 1998). Progeny transgenic seeds were screened on hygromycin until the T_4_ generation. For live cell F-actin imaging, wild type and *lbr1* were transformed with the *UBQ10::GFP-ABD2-GFP* construct (Dyachok *et al*., 2014).

### Statistical analysis

One-way analysis of variance (ANOVA) tests and Tukey’s multiple mean comparisons were performed using the lsmeans package (Lenth, 2016) in R (R Core Team, 2019). Box plots were generated with the ggplot function in the ggplot2 package (Wickham, 2016).

## Results

### Isolation and characterization of an *Arabidopsis thaliana* mutant that is resistant to latrunculin B

A forward genetic screen was performed with EMS-mutagenized *A. thaliana* Col-0 ecotype seeds grown on LatB to identify new regulators of actin-mediated root development. The primary root growth of wild type seedlings is severely inhibited by 100 nM LatB (Fig. 1A, B), allowing for a convenient forward genetic screen for mutants that are resistant to this compound. From this screen, we identified the mutant *lbr1*.

**Figure 1.**
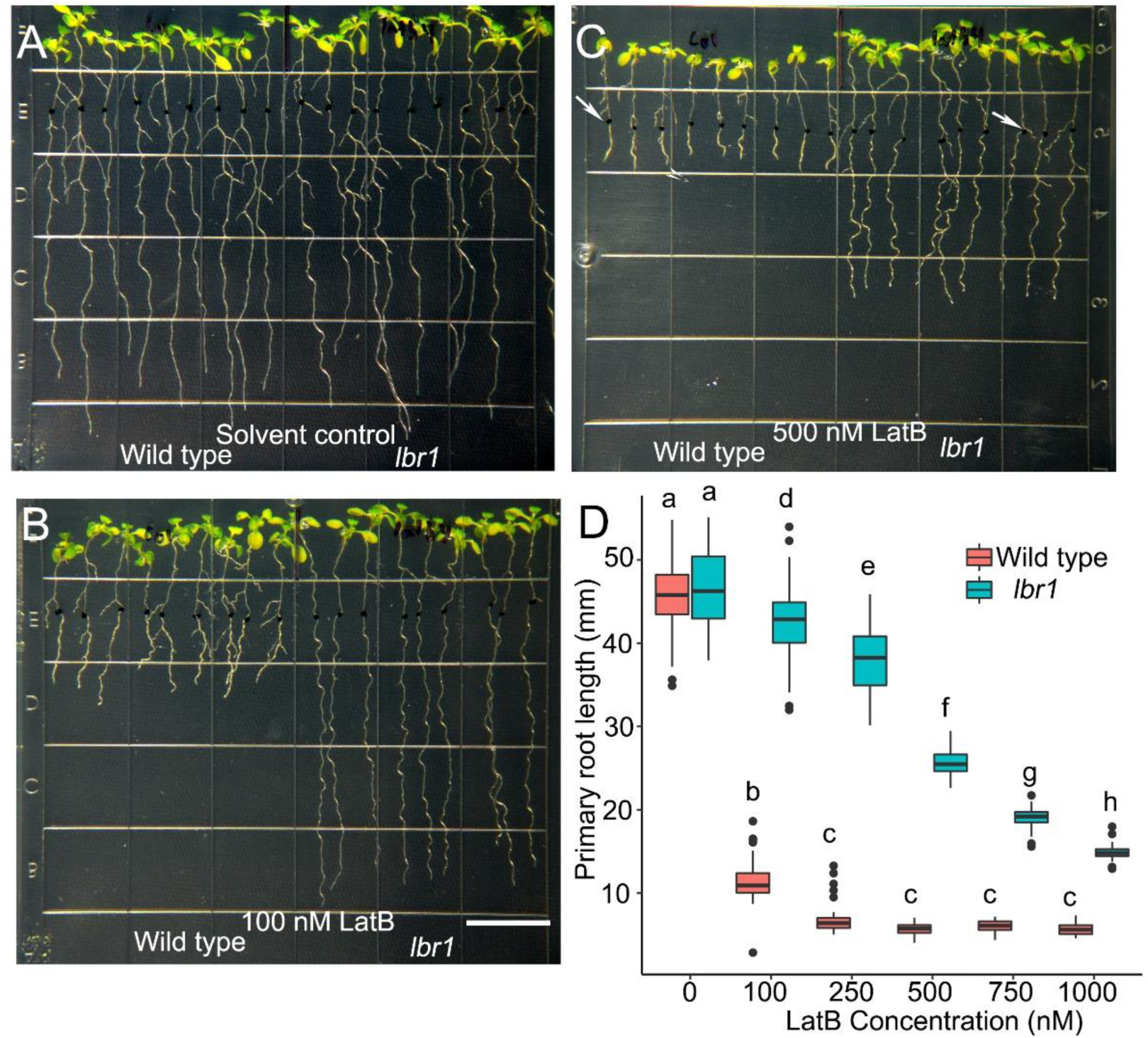
Primary roots of *lbr1* are more resistant to the growth inhibitory effects of LatB than those of wild type. Representative images of three-day-old seedlings of wild type and *lbr1* were transplanted to medium without LatB (solvent controls, **A**), and supplemented with 100 nM **(B)**, or 500 nM LatB **(C),** and grown for an additional six days. Black dots (e.g., white arrows in **C**) in **A**-**C** indicate the position of the root tip upon transplant. Bar in **B** is 13 mm. (**D**) Box plots of primary root elongation of wild-type and *lbr1* seedlings on solvent control (0) and various LatB concentrations. Box limits indicate 25^th^ and 75^th^ percentiles, the horizontal line is the median, and whiskers display minimum and maximum values. Quantification of the primary root length was obtained by measuring the distance between the root tip after six days and the black dot. Statistical significance in **D** was determined by one-way ANOVA. *n* = 50-60 seedlings. Different letters are significantly different (*p* < 0.05, Tukey’s test).

There were no statistically significant differences in the elongation of wild type and *lbr1* primary roots on medium without LatB (Fig. 1A, D). By contrast, primary roots of 3-day-old wild type seedlings transferred to medium containing 100 nM LatB were severely inhibited six days later (i.e., roots showed >75% growth inhibition), while those of *lbr1* were only partially inhibited (Fig. 1B, D). The longer roots of *lbr1* on LatB relative to wild type was also evident at concentrations of 500 and 1000 nM (Fig. 1C, D).

Tolerance of *lbr1* to LatB was also manifested in other aspects of seedling development. For example, *A. thaliana* seedlings in complete darkness exhibited the typical elongated hypocotyl response. Without LatB in the medium, wild type and *lbr1* seedlings had similar hypocotyl lengths in the dark (Supplementary Fig. S2A, C). By contrast, when seedlings were grown in the dark on medium containing 100 nM or 500 nM LatB, wild type hypocotyls were significantly shorter than those of *lbr1* (Supplementary Fig. S2B, C).

We next analyzed root hair length of *lbr1* on LatB given the known role of actin in tip growth (Bascom *et al*., 2018). On growth medium without LatB, *lbr1* root hair length was slightly longer than that of wild type. Differences between *lbr1* and wild type root hairs became more pronounced when seeds were germinated directly in low concentrations of LatB with *lbr1* showing longer root hairs than that of wild type (Supplementary Fig. S2D, E).

### A point mutation in *ACTIN7* is responsible for latrunculin B resistance in *lbr1*

The *lbr1* mutation was backcrossed six times to remove extraneous, unlinked mutations. During backcrossing, root phenotypes of the F_1_ and F_2_ progenies on 100 nM LatB were evaluated to determine whether *lbr1* is dominant or recessive. Results revealed that 100% of the F_1_ seedlings were resistant to 100 nM LatB, while the segregating F_2_ progenies showed a 3:1 LatB resistant: LatB sensitive ratio (Supplementary Fig. S3A, B). These results indicate that *lbr1* is a dominant-negative mutant.

Map-based cloning combined with next-generation sequencing revealed that *lbr1* had a single nucleotide mutation in the *AT5G09810* gene, which encodes the vegetative actin isoform ACT7. The mutation in the *ACT7* coding sequence of *lbr1* changed the cytosine at position 100 located in the 2^nd^ exon to a thymine, leading to a substitution of proline-34 (pro-34) with serine (ACT7_P34S_; Supplementary Fig. S3C). ACT7_P34S_ is located adjacent to the nucleotide-binding cleft of the actin protein monomer (Supplementary Fig. S3D; Morton *et al*., 2000; Iwabuchi *et al*., 2019).

Evidence confirming that the ACT7 pro-34 to ser change is responsible for the *lbr1* phenotype was obtained by introducing *ACT7_P34S_* under the control of the *ACT7* promoter (*ACT7pro*:*ACT7_P34S_*) into wild type. When transgenic seedings were grown on 100 nM LatB, they showed resistance to LatB that was similar to that of *lbr1* (Fig. 2A, B). Further validation that *lbr1* mutation was due to the pro-34 to ser change in *ACT7*, wild type *ACT7* (*ACT_WT_*) under the control of *ACTpro* (*ACT7pro*:*ACT7_WT_*) was introduced to *lbr1*. *lbr1* transformed with *ACT7pro*:*ACT7_WT_* exhibited wild type sensitivity to LatB (Fig. 2C, D).

**Figure 2.**
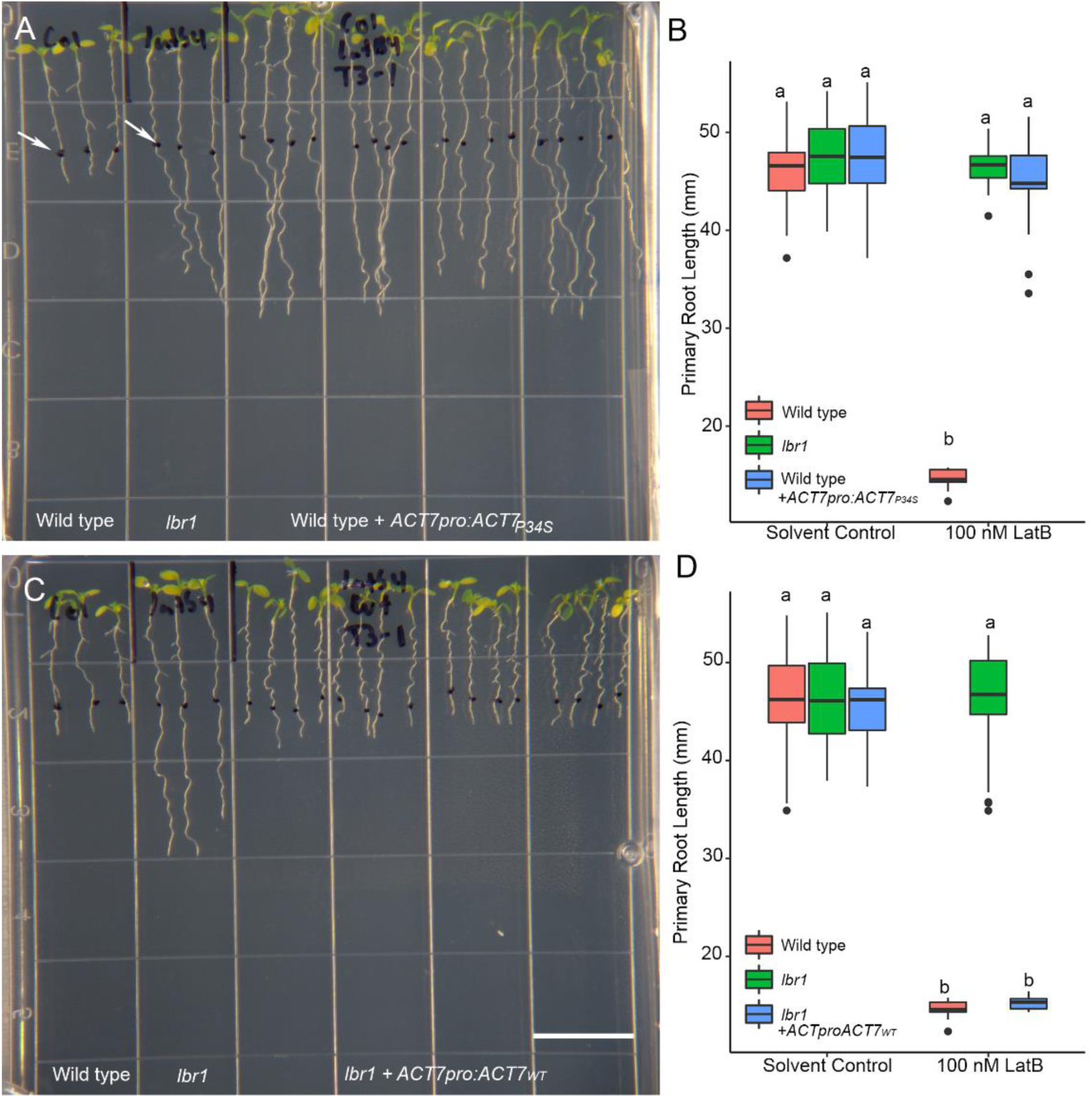
Transgenic complementation of the *lbr1* mutant confirms that proline to serine substitution in position 34 (P34S) in ACT7 leads to LatB resistance. (**A, B**) Wild type seedlings transformed with the *ACT7pro*:*ACT7_P34S_* are as resistant to LatB as *lbr1*. Bar in **B** = 13 mm. (**C, D**) *lbr1* seedlings lose their resistance to LatB when transformed with the *ACT7pro*:*ACT7_WT_* construct. Box limits in **B** and **C** indicate 25^th^ and 75^th^ percentiles, horizontal line is the median, and whiskers display minimum and maximum values. Quantification of the primary root length was obtained by measuring the distance between the root tip after six days and the black dot, which marks the position of the root tip during seedling transplant (white arrows in **A**). Statistical significance was determined by one-way ANOVA. *n* = 25-35 seedlings. Different letters are significantly different (*p* < 0.05, Tukey’s test).

Cytochalasins are another class of inhibitors used to study actin function (Schliwa,1982). We therefore asked if *lbr1* also exhibits resistance to cytochalasin D. We found that *lbr1* showed wild-type sensitivity to cytochalasin D, indicating that the *ACT7_P34S_* mutation is specific to LatB (Supplementary Fig. S4).

### Latrunculin B disrupts filamentous actin in *lbr1* roots to a lesser extent than that of wild type

We hypothesized that filamentous-actin (F-actin) in the *lbr1* primary roots is affected to a lesser extent by LatB than that of wild type given the root phenotypes described above. This hypothesis was tested by characterizing roots of wild type and *lbr1,* both expressing the live F-actin reporter *UBQ10:GFP-ABD2-GFP* (Dyachok *et al*., 2014). The overall appearance of F-actin in epidermal cells in the root elongation zone of wild type was similar to that of *lbr1* seedlings grown in solvent control media for six days (Fig. 3A). On the other hand, growth on 100 nM LatB for six days led to a visible reduction in the F-actin density in epidermal cells of the root elongation zone in wild type seedlings, but not in *lbr1* (Fig. 3B, C). Consistent with visual observations, quantitative analysis showed that the F-actin density in the cells of the root elongation zone of untreated wild type seedlings was similar to that of *lbr1*. By contrast, although F-actin density in roots of *lbr1* treated with 100 nM LatB was significantly less than that of untreated plants, it was significantly higher than wild type treated with 100 nM LatB (Fig. 3D), consistent with *lbr1* being resistant to the effects of LatB on actin structure.

**Figure 3.**
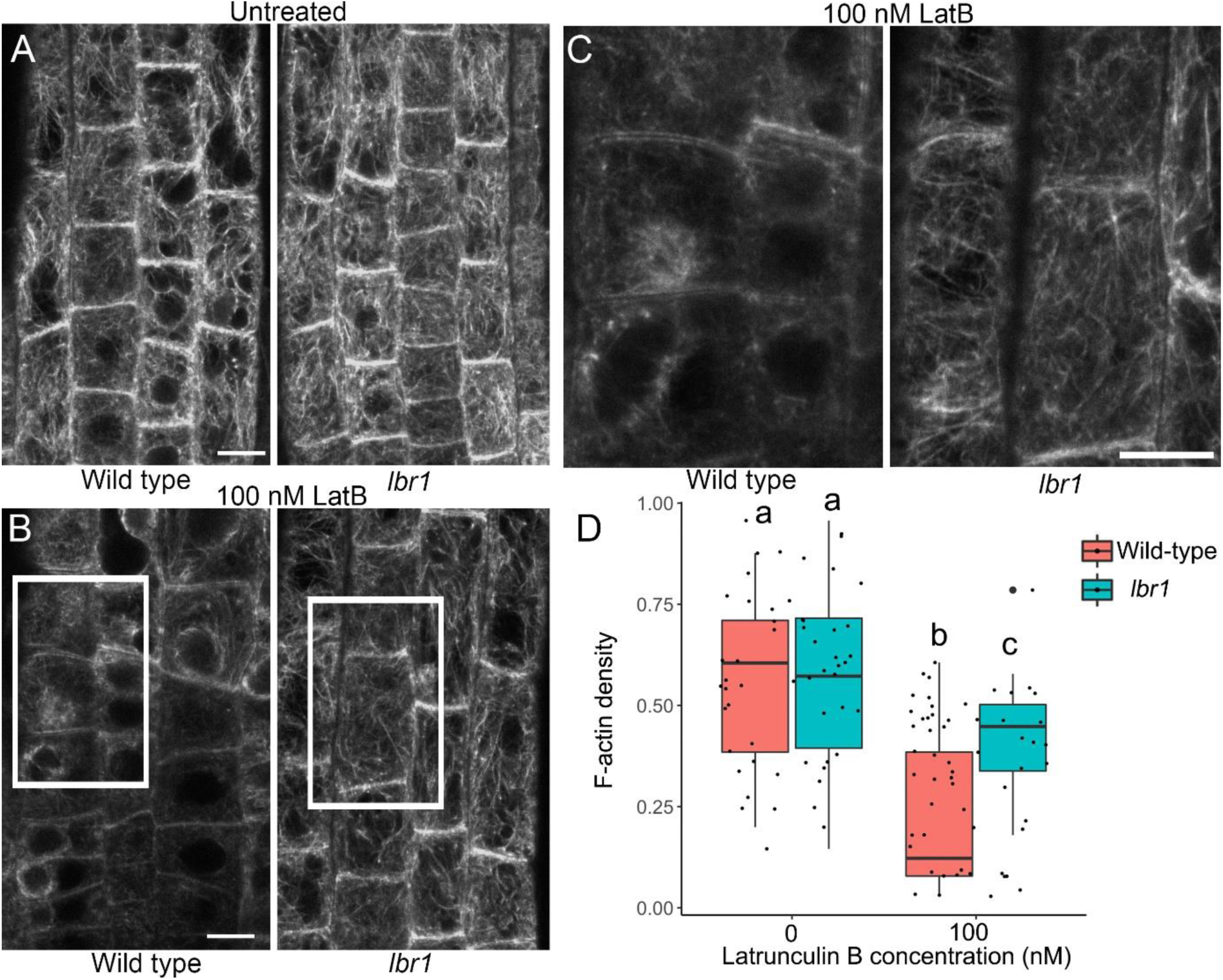
Density of F-actin in epidermal cells in the elongation zone of *lbr1* primary roots is higher than that of wild type when exposed to LatB. Representative confocal micrographs of GFP-labeled F-actin in untreated (**A**) and 100 nM LatB (**B**) wild type and *lbr1* roots. (**C**) Higher magnification images of cells corresponding to the white rectangles in **B**. Higher magnification images were used to quantify F-actin density as described in Supplementary Figure S1. (**D**) Quantification of F-actin density in root epidermal cells. Box limits indicate 25^th^ and 75^th^ percentiles, horizontal line is the median, and whiskers display minimum and maximum values. Statistical significance was determined by one-way ANOVA. *n* = 24-32 images. Different letters are significantly different (*p* < 0.05, Tukey’s test). Bars in **A**-**C** = 20 μm.

### *ACT7_P34S_* mutation confers partial latrunculin B resistance to an *ACT7* mutant with severe developmental defects

We next asked if the *ACT7_P34S_* mutation confers LatB resistance to an *ACT7* mutant that exhibits severe seedling developmental defects. One such mutant is a*ct7-5*, which is a recessive null mutant allele with a T-DNA insertion in the third exon of the *ACT7* gene (Kandasamy *et al*., 2010). We found that the *ACT7pro*:*ACT7_P34S_* complemented the severe seedling developmental defects of *act7-5*, indicating that *ACT7_P34S_* encodes a functional ACT7 variant (Fig. 4A, C). Importantly, seedling roots of *act7-5* complemented with *ACT7pro*:*ACT7_P34S_* exhibited resistance to LatB (Fig. 4B,C). However, LatB resistance of *act7-5* primary roots complemented with *ACT7pro*:*ACT7_P34S_* was not as strong as that of *lbr1* or wild-type seedlings expressing *ACT7pro*:*ACT7_P34S_* (Fig. 4B, C) On the other hand, the hypocotyl length of dark-grown *act7-5* complemented with *ACT7pro*:*ACT7_P34S_* grown on LatB was similar to that of wild type (Fig. 4D).

**Figure 4.**
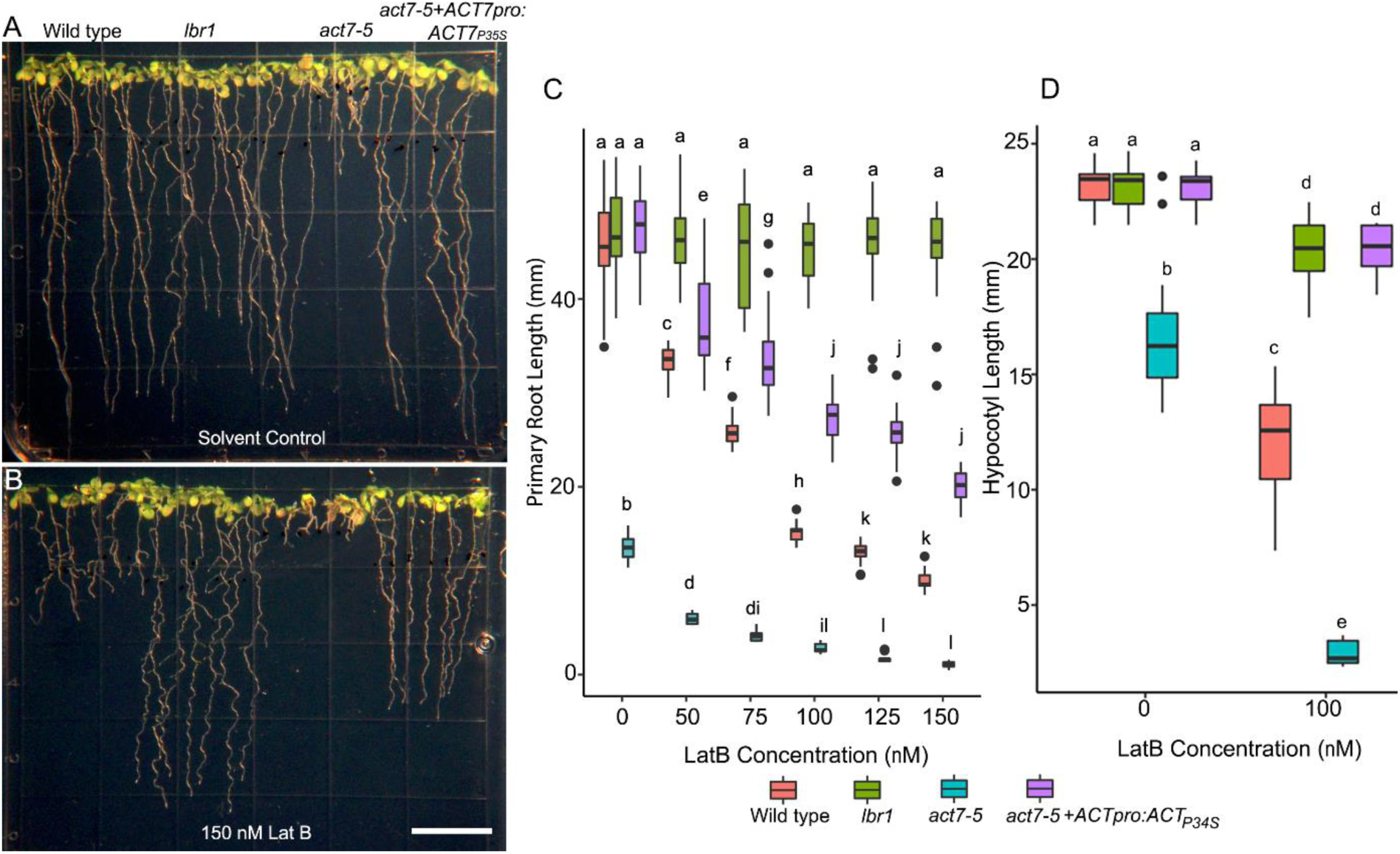
The ACT7 P34S mutation confers partial LatB resistance to *act7-5*. Representative images of seedlings of wild type, *lbr1*, *act7-5*, and *act7-5* transformed with *ACT7pro:ACT7_P34S_* in solvent controls (**A**) or 150 nM LatB (**B**). Quantification of primary root length of light-grown seedlings (**C**) and hypocotyl length in dark grown (**D**) wild type, *lbr1*, *act7-5*, and *act7-5* transformed with *ACT7pro:ACT7_P34S_* seedlings with and without LatB. Box limits indicate 25^th^and 75^th^ percentiles, horizontal line is the median, and whiskers display minimum and maximum values. Statistical significance was determined by one-way ANOVA. *n* = 25-35 seedlings. Different letters are significantly different (*p* < 0.05, Tukey’s test).

### A P34S mutation in ACT2 confers latrunculin B resistance

Given the 94 – 95% amino acid sequence similarity between ACT2 and ACT7, we next asked if changing the amino acid proline at position 34 to a serine in the *ACT2* gene could also lead to LatB resistant seedlings. To address this question, we performed site-directed mutagenesis in the wild-type *ACT2* cDNA, in which the cytosine at position 100 in the *ACT2* coding sequence was changed to a thymine. This mutation led to a substitution of serine for proline in ACT2 that mirrored that of LBR1/ACT7_P34S_ (see Supplementary Fig. S3). We designated this *ACT2* point mutant as *ACT2_P34S_*. The *ACT2_P34S_* gene was then cloned into the *pCAMBIA1390* vector with the *ACT2* promoter to generate a *ACT2pro:ACT2_P34S_* construct. As a control, the wild type *ACT2* gene was also cloned into the *pCAMBIA1390* vector to generate the *ACT2pro:ACT2_WT_* construct. Results showed that primary roots and dark-grown hypocotyls of wild type and *ACT2pro:ACT2_WT_* seedlings exhibited similar sensitivity to LatB (Fig. 5A, C). Like *lbr1*, *ACT2pro:ACT2_P34S_* seedlings were more resistant to LatB than wild type seedlings (Fig. 5B, D). However, the resistance of *ACT2pro:ACT2_P34S_* seedling roots to LatB was less than that of *lbr1* (Supplementary Fig. S5).

**Figure 5.**
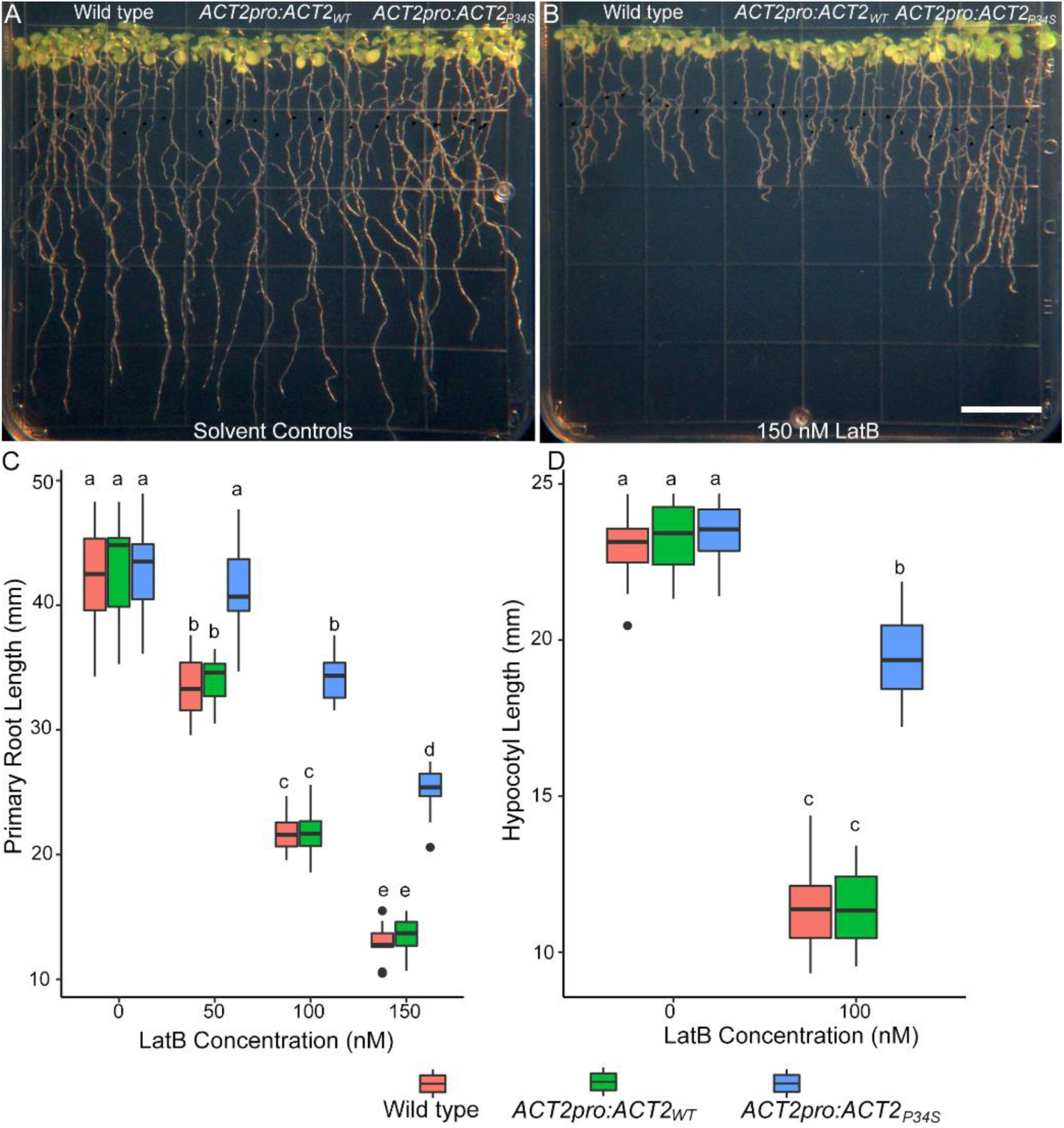
The ACT2 P34S mutation confers partial LatB resistance to wild type seedlings. Representative images of wild type and wild type transformed with *ACT2pro:ACT2_WT_* and *ACT2pro:ACT2_P34S_* in solvent controls (**A**) or 150 nM LatB (**B**). Note that *ACT2pro:ACT2_P34S_*, but not *ACT2pro:ACT2_WT_* confers LatB resistance. Bar in **B** = 13 mm. Quantification of primary root length of light-grown seedlings (**C**) and hypocotyl length in dark-grown (**D**) wild type and wild type transformed with *ACT2pro:ACT2_WT_* or *ACT2pro:ACT2_P34S_* with or without LatB. Box limits indicate 25^th^ and 75^th^ percentiles, horizontal line is the median, and whiskers display minimum and maximum values. Statistical significance was determined by one-way ANOVA. *n* = 25-35 seedlings. Different letters are significantly different (*p* < 0.05, Tukey’s test).

### Targeted mutagenesis of ACT7 reveals amino acid residues critical for latrunculin B effects on root development

As noted, the pro-34 to ser mutation in ACT7 of *lbr1* is located adjacent to the nucleotide-binding cleft of the G-actin monomer (Supplementary Fig. S2D). Earlier studies in yeast showed that mutations close to the nucleotide-binding cleft of the actin protein confer resistance to latrunculin A (LatA), which is structurally similar to LatB (Ayscough *et al*., 1997). Furthermore, synthetic *O*-methylated or *N*-alkylated LatB analogues that lose their ability to form hydrogen bonds with amino acid residues adjacent to the actin nucleotide-binding cleft have low bioactivity (Kudrimoti *et al*., 2009). We therefore hypothesized that mutations in other amino acid residues in equivalent regions adjacent to the *A. thaliana* ACT7 nucleotide-binding cleft that form hydrogen bonds with LatB will result in LatB-resistant seedlings that mirror those of *lbr1*. To test this hypothesis, five amino acid residues in the actin protein that form hydrogen bonds with latrunculin based on synthetic LatB analogue docking studies and analyses of the crystal structures of actin-latrunculin complexes were selected. These included the amino acid residues Tyrosine (Y)-69, Aspartate (D)-157, Threonine (T)-186, Glutamine(E)-207, and Arginine (R)-210 (Morton *et al*., 2000; Kudrimoti *et al*., 2009). Sequence alignment with mammalian and yeast actin indicates that the equivalent *A. thaliana* ACT7 amino acid residues are Tyrosine-71(Y71), Aspartate-159 (D159), Threonine-188 (T188), Glutamine-209 (E209), and Arginine-211 (R211), respectively (Fig. 6A). The Y71, D159, T186, E209, and R211 residues were replaced with glutamate (G) to generate the site-directed ACT7 mutants Y71G, D159G, T186G, E209G, and R211G ACT7. Mutants were separately cloned into the *pCAMBIA1390* vector under the control of the *ACT7* promoter and transformed into wild type to generate *ACT7pro:ACT7_Y71G_, ACT7pro:ACT7_D159G_, ACT7pro:ACT7_T188G_*, *ACT7pro:ACT7_E209G_*, and *ACT7pro:ACT7_R211G_* lines.

**Figure 6.**
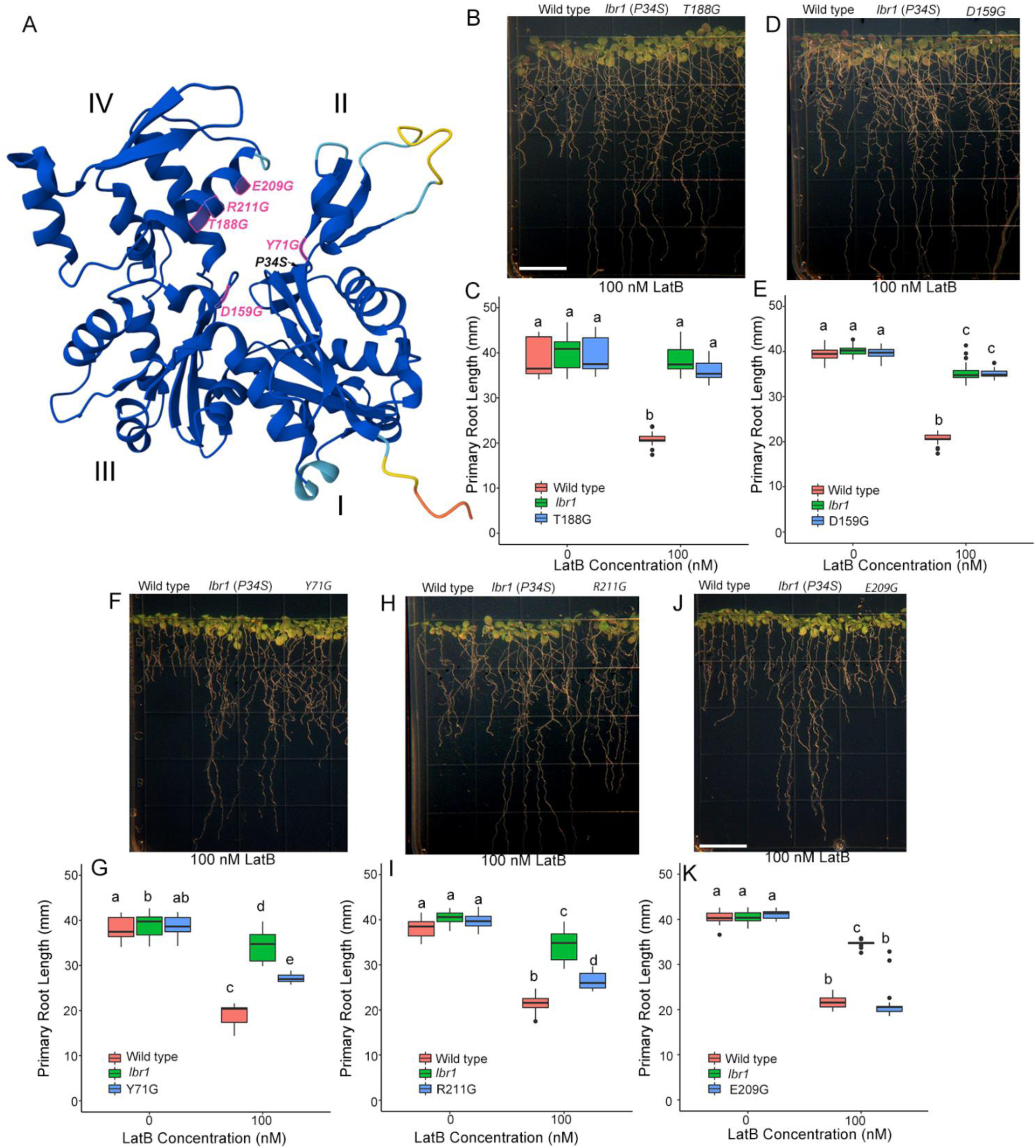
Site-directed mutagenesis reveals amino acid residues within the ACT7 nucleotide-binding cleft that are crucial for conferring LatB resistance. (**A**) Ribbon model of the ACT7 protein monomer showing the location of P34S mutation and five site-directed mutants (in magenta text). The ribbon structure and location of site-directed mutants were obtained using AlphaFold version 2 (Skolnick *et al*., 2021) using the amino acid sequence of the *A. thaliana* ACT7. Roman numerals indicate the four domains of the actin monomer. (**B-K**) Representative seedling images and quantification of primary root elongation of site-directed ACT7 mutants Y71G, D159G, T186G, E209G, and R211G ACT7. Box limits indicate 25^th^ and 75^th^ percentiles, horizontal line is the median, and whiskers display minimum and maximum values. Statistical significance was determined by one-way ANOVA. *n* = 25-35 seedlings. Different letters are significantly different (*p* < 0.05, Tukey’s test). Bars in **B** and **J** = 13 mm.

Primary root growth of seedlings expressing the five site-directed ACT7 mutants was evaluated by growing them in LatB-free medium for 3 days and then transferring them to solvent control or 100 nM LatB medium and grown for an additional six days as described above. We found that roots of *ACT7pro:ACT7_D159G_ and ACT7pro:ACT7_T188G_* lines exhibited similar resistance to 100 nM LatB as that of *lbr1* (Fig. 6B-E). Roots of seedlings expressing *ACT7pro:ACT7_Y71G_* and *ACT7pro:ACT7_R211G_* also exhibited resistance to 100 nM LatB but such resistance was less than that of *lbr1* (Fig. 6F-I). By contrast, roots of seedlings expressing *ACT7pro:ACT7_E209G_* showed similar sensitivity to 100 nM LatB as that of wild type (Fig. 6J, K).

## Discussion

Latrunculins are metabolites isolated from the red sea sponge *N. magnifica* capable of rapidly and reversibly disrupting the actin cytoskeleton in a variety of eukaryotic cells, making them valuable pharmacological tools to study actin function (Spector *et al*., 1983; Morton *et al*., 2000). The use of LatB has led to insights into the roles of actin in a number of plant biological processes, including gravitropism, tip growth, root and trichome development, and plant-microbe interactions (Mathur *et al*., 1999; Gibbon *et al*., 1999; Baluška *et al*., 2001b; Blancaflor, 2013; Leontovyčová *et al*., 2020; Hiles *et al*., 2024).The distinct inhibitory effects of LatB on primary root growth are of particular interest because this has paved the way for forward genetic screens that are instrumental in identifying new molecular components that modulate actin function in plants. For example, a class of recessive *A. thaliana* mutants hypersensitive to LatB have uncovered proteins that link the actin cytoskeleton to membrane trafficking and the actin-nucleating machinery modulated by the SCAR-WAVE complex (Sparks *et al*., 2016; Chin *et al*., 2021). Here, we describe the characterization of the *A. thaliana lbr1* mutant that is resistant to LatB. Genetic analyses showed that *lbr1* is a new dominant-negative mutant of *ACT7*, which encodes one of three vegetative ACT isoforms in plants (Kandasamy *et al*., 2010). Our work on *lbr1* described here provides insights into LatB’s mechanism of action on plant actin that appears to be mostly conserved with that of other eukaryotic actins.

Map-based cloning and whole genome sequencing revealed that *lbr1* has a mutation in the proline residue at position 34 (P34), which is located in domain I of the actin monomer (Supplementary Fig. S2). This proline residue clusters around a region adjacent to the actin nucleotide-binding site based on previous LatA-resistant yeast mutant and actin-LatA crystallography studies (Ayscough *et al*., 1997; Morton *et al*., 2000). The R183 and D184 residues also cluster to a region adjacent to the actin nucleotide-binding cleft, and when mutated, lead to LatA resistant yeast cells. Corresponding R183 and D184 mutants were also able to confer LatA resistance in mammalian HeLa cells, indicating that the mechanism of action of latrunculins, in which the molecule interferes with nucleotide exchange to prevent actin polymerization, is conserved in yeast and mammals (Morton *et al*., 2000; Fujita *et al*., 2003). However, it is unknown to what extent latrunculin’s mechanism of action extends to plant actin. The location of ACT7_P34S_ mutation indicates that plant actin shares similar features as yeast and mammalian actin. To further test this hypothesis, we generated five site-directed ACT7 mutants that corresponded to the amino acid residues mutated in yeast LatA resistant mutants and expressed them in *A. thaliana* (Ayscough *et al*., 1997; Morton *et al*., 2000). We found that four of the five site-directed mutants in ACT7 conferred LatB resistance. ACT7_T188G_ and ACT7_D159G_ led to LatB resistance that is comparable to that of ACT7_P34S_. ACT7_Y71G_ and ACT7_R211G_ also conferred LatB resistance but were not as strong as that of ACT7_P34S_, ACT7_T188G_, and ACT7_D159G_ (Fig. 6). These results indicate that the latrunculin binding site reported in yeast is conserved in plant vegetative actin. For instance, the D157 and T186 positions in subdomain III and IV of yeast actin, respectively, form hydrogen bonds with latrunculin (Morton *et al*., 2000). The corresponding ACT7_D159G_ and ACT7_T188G_ conferred strong LatB resistance in *A. thaliana* (Fig. 6). These amino acid residues are directly above the nucleotide-binding cleft of the yeast actin and *A. thaliana* ACT7 monomer (Morton *et al*., 2000; Fig. 6). The closer proximity of ACT7_P34S_, ACT7_T188G_, and ACT7_D159G_ to the nucleotide-binding cleft of ACT7 relative to ACT7_Y71G_, and ACT7_R211G_ could explain why expression of the latter leads to a more robust LatB resistant phenotype (Fig. 6). E207 in subdomain IV also forms a hydrogen bond with LatA based on computational docking and semisynthetic LatA analog studies (Kudrimoti *et al*., 2009). To the best of our knowledge, however, there are no yeast or mammalian mutants to E207. As such, evidence implicating this residue in latrunculin bioactivity is lacking. We therefore generated an ACT7_E209G_ mutant that corresponds to E207 and found that it does not confer resistance to LatB. This result suggests minimal contribution of the E207, E209 residues to LatB’s mechanism of action.

There are three other dominant-negative mutants identified in *A. thaliana* vegetative actin isoforms. The *act2-2D* and *fryzzy1actin8D*, which have mutations in *ACT2* and *ACT8*, respectively, display severe plant developmental and actin organization defects (Nishimura *et al*., 2003; Kato *et al*., 2010). More recently, a mutant called *unp1*, which exhibits abnormal dark-induced nuclear positioning and fragmented F-actin in leaves, was shown to contain a mutation in the ACT7 DNAse I-binding loop (D-loop) (Iwabuchi *et al*., 2019). Interestingly, expression of *unp1*-like mutations to *ACT2* and *ACT8* resulted in similar dark-induced nuclear positioning defects as those of *unp-1*, implicating involvement of all three vegetative actins in this process. Our results that a *lbr1*-like, P34S mutation in *ACT2* confers LatB resistance support previous studies demonstrating overlapping functions of actin isoforms in plants (Kandasamy *et al*., 2010) and that LatB effects are facilitated through the same region within the vegetative actin isoforms. The ability of ACT7_P34S_ to complement the severe developmental defects of *act7-5* and the lack of F-actin and seedling phenotypes in *lbr1* indicate that the mutation does not significantly affect ACT7’s core functions. Based on the results presented here, P34S and adjacent amino acid residues affect mostly the interaction between LatB and ACT7. The observation that *lbr1* is not resistant to cytochalasin D further supports specificity to LatB. It would be interesting to test whether *unp1* shows differential sensitivity to LatB because of the crucial role of the D-loop in actin polymerization (Baek *et al*., 2008).

In conclusion, the isolation and characterization of *lbr1* further illustrate the power of forward genetic screens using LatB to elucidate the role of the actin cytoskeleton in plant development as we have previously shown (Sparks *et al*., 2016; Sun *et al*., 2019; Chin *et al*., 2021). Although the manner by which latrunculin binds to G-actin is known in yeast and mammals, it remains unclear whether similar mechanisms extend to plant G-actin. Our studies of the dominant-negative *lbr1* mutant now indicate that latrunculin’s mechanism of action among eukaryotic actins is conserved. The set of ACT7 amino acid mutants generated here also paves the way for studies to probe deeper into the molecular nature of actin function in plant cells. For example, given that some actin regulatory proteins, such as profilin and gelsolin, modulate actin polymerization through nucleotide exchange (Morton *et al*., 2000), it would be interesting to ask whether LatB-resistant site-directed mutants impact the binding and activity of plant actin regulatory proteins. Finally, the discovery of LatB-resistant mutants holds significant potential for developing plants with modified responses to environmental stresses, particularly those that exert their impact on plant development through the actin cytoskeleton.

## Supplementary Data

The following supplementary data are available at JXB online:

**Fig. S1.**
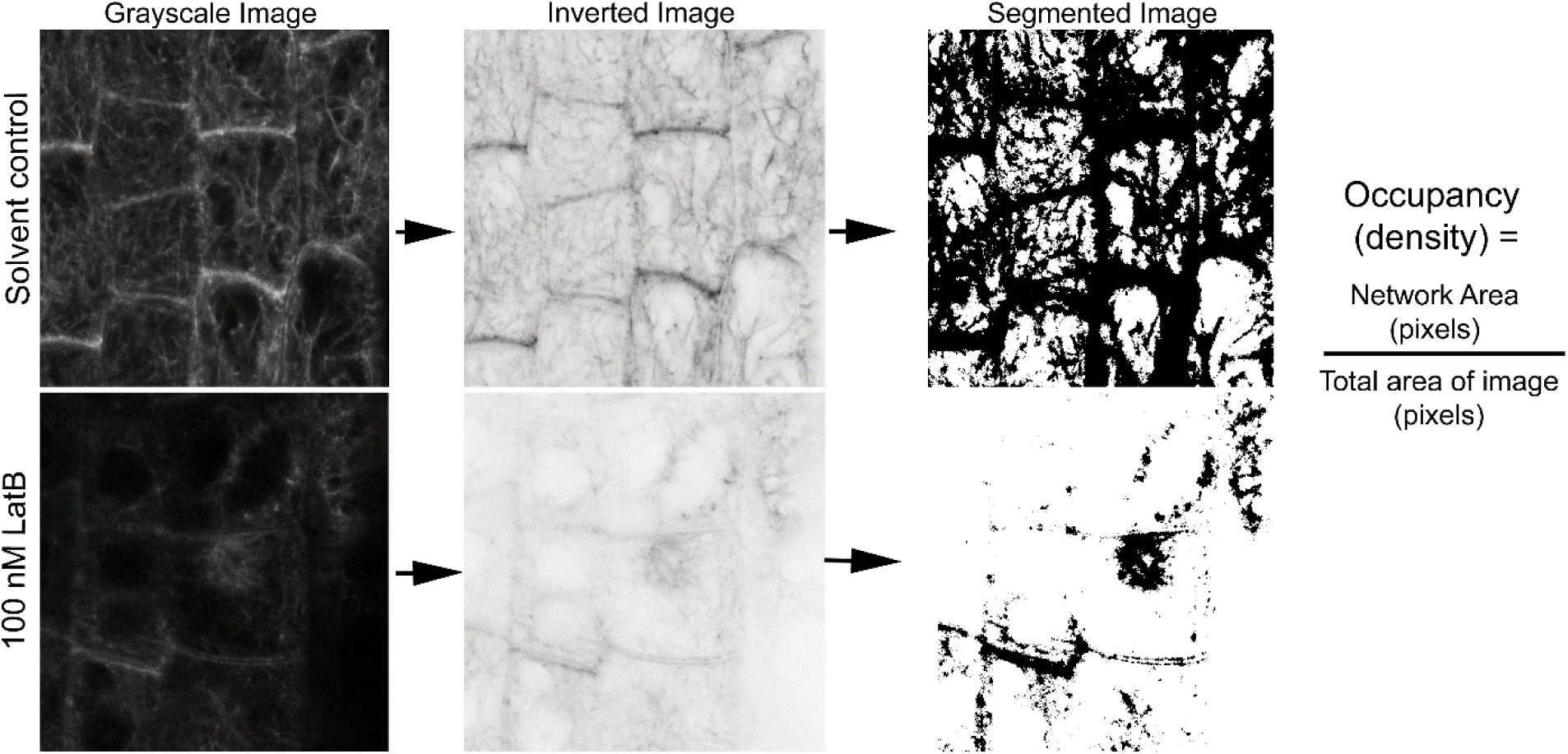
Overview of F-actin density quantification in root epidermal cells using RhizoVision. Greyscale confocal micrographs in TIFF format were first inverted using Photoshop to obtain black filaments in white background. Inverted images were then dragged and dropped in RhizoVision Explorer v2.0.3 (https://doi.org/10.5281/zenodo.3747697; <u>Seethepalli and York</u> <u>2020</u>) through the file menu. Inverted images were segmented using the same threshold settings for solvent controls and LatB-treatment using the analyze menu. The area in black pixels, which correspond to the F-actin network, was divided by total pixels in the entire image to obtain density.

**Fig.S2.**
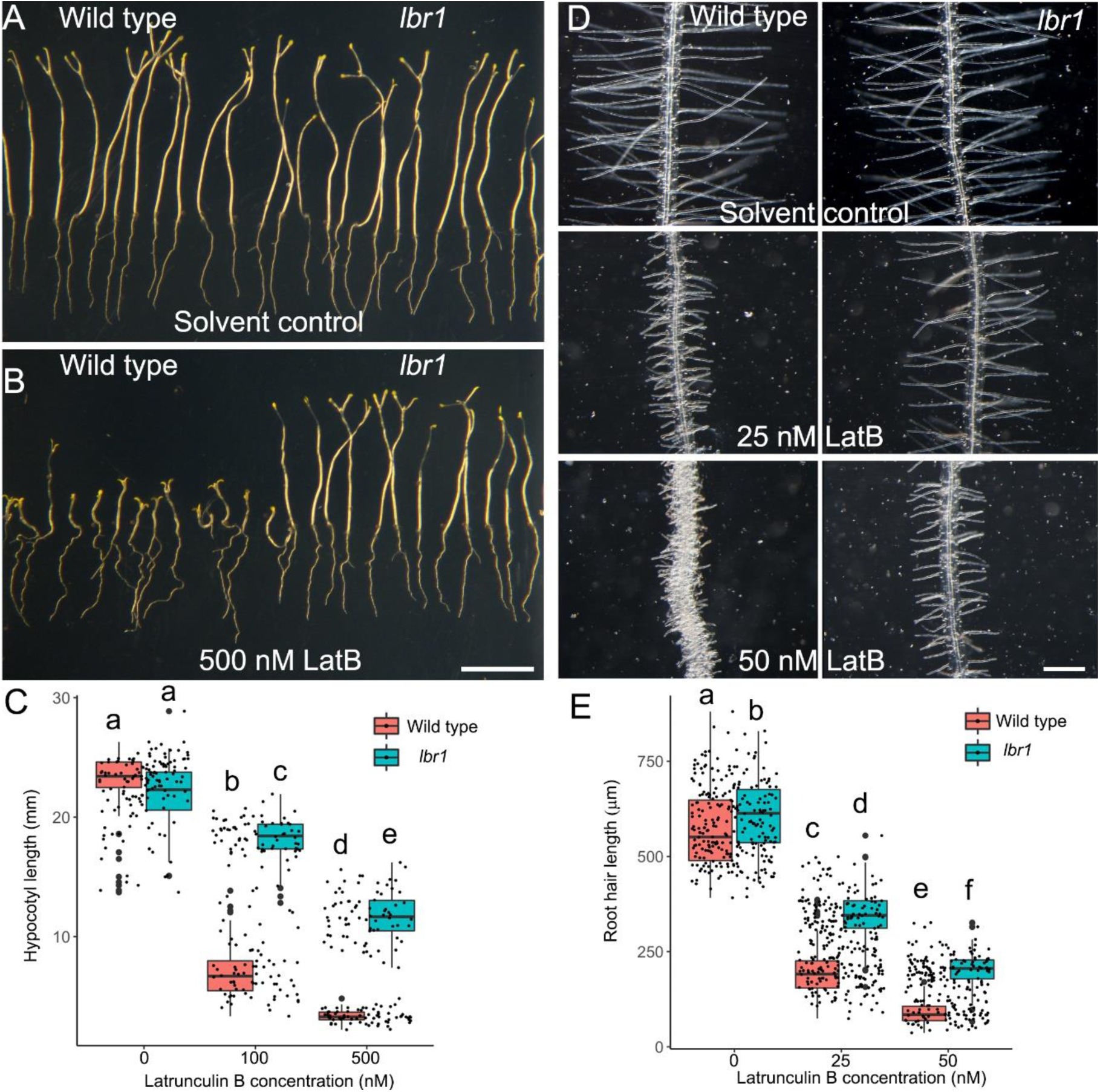
LatB resistance of *lbr1* is manifested in other aspects of seedling development. Representative images of wild-type and *lbr1* seedlings transplanted to medium without LatB (solvent controls, **A**) or supplemented with 500 nM **(B)** and kept in total darkness for five days. (**C**) Box plots of hypocotyl length of wild-type and *lbr1* seedlings on 0 (solvent control), 100 and 500 nM LatB. (**D**) Representative root hair images of wild-type and *lbr1* primary roots from seeds directly germinated media without LatB (solvent controls) or supplemented with 25 or 50 nM. (**E**) Box plots of root hair length of wild-type and *lbr1* seedlings on 0 (solvent control), 25 and 50 nM LatB. Box limits indicate 25^th^ and 75^th^ percentiles, horizontal line is the median, and whiskers display minimum and maximum values. Statistical significance in **C** and **E** was determined by one-way ANOVA. *n* = 50-60. Different letters are significantly different (*p* < 0.05, Tukey’s test). Bar in **B** for **A** and **B** = 10 mm; Bar in **D** = 200 μm.

**Fig. S3.**
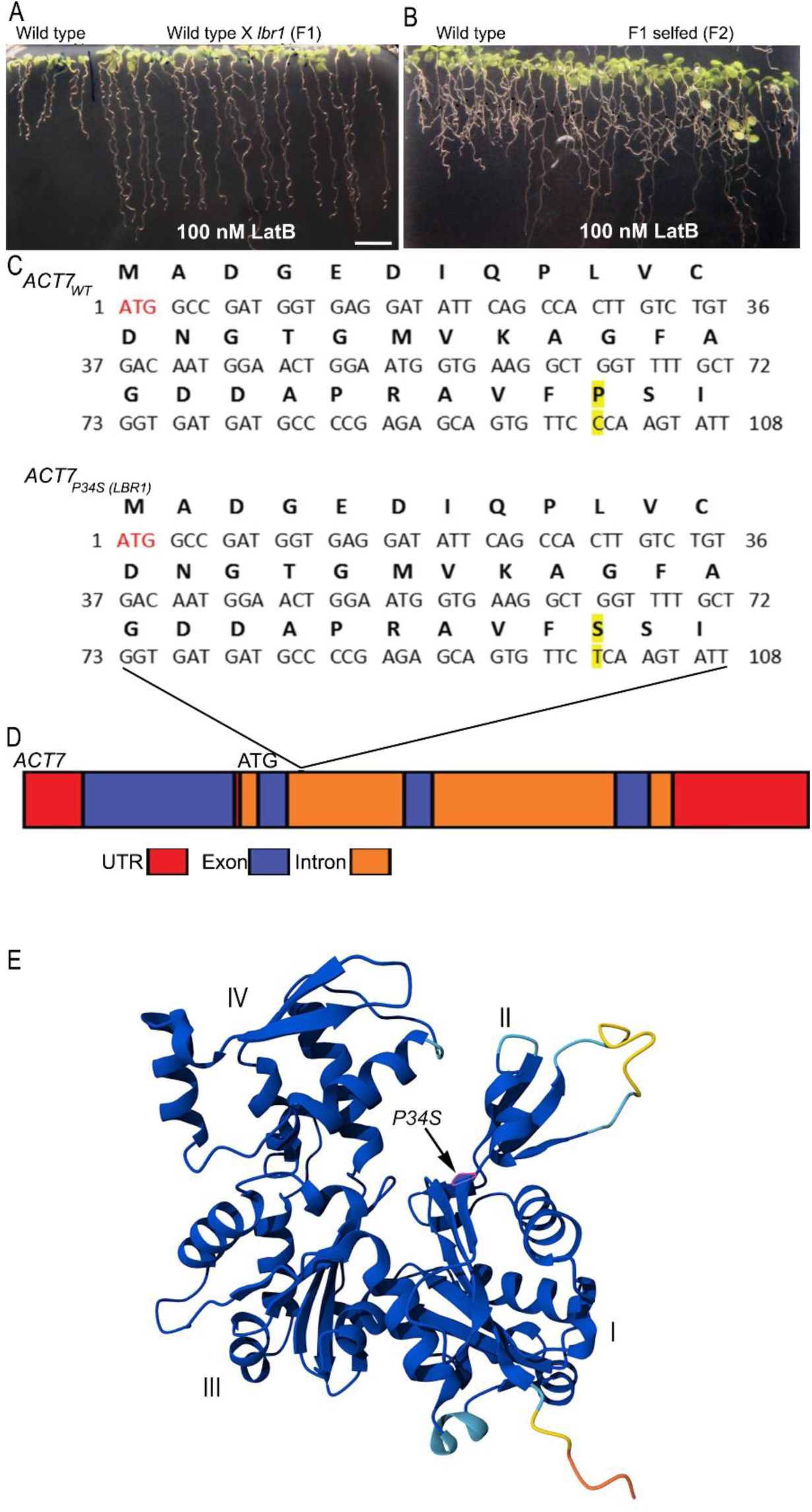
Genetic characterization and cloning of *LBR1*. (**A**) A cross between wild type and *lbr1* resulted in F1 progeny that was resistant to LatB. (**B**) The F2 population from selfed F1 led to a ration of 3 LatB resistant to 1 wild type. (**C**) Nucleotide and amino acid sequence of *ACT7_WT_* and *ACT7_LBR1_* showing the position of the cytosine (C) to thymine (T) mutation (yellow highlights) that led to a proline to serine substitution at position 34 of ACT7 in the *lbr1* mutant. (**D**) Genome organization of *ACT7* showing that the C to T mutation is located in the second exon. (**E**) Ribbon model of the ACT7 protein monomer based on alphafold 2 (Skolnick et. al., 2021) showing the location of P34S mutation adjacent to the nucleotide-binding cleft. Numbers indicated four domains of the actin monomer.

**Fig. S4.**
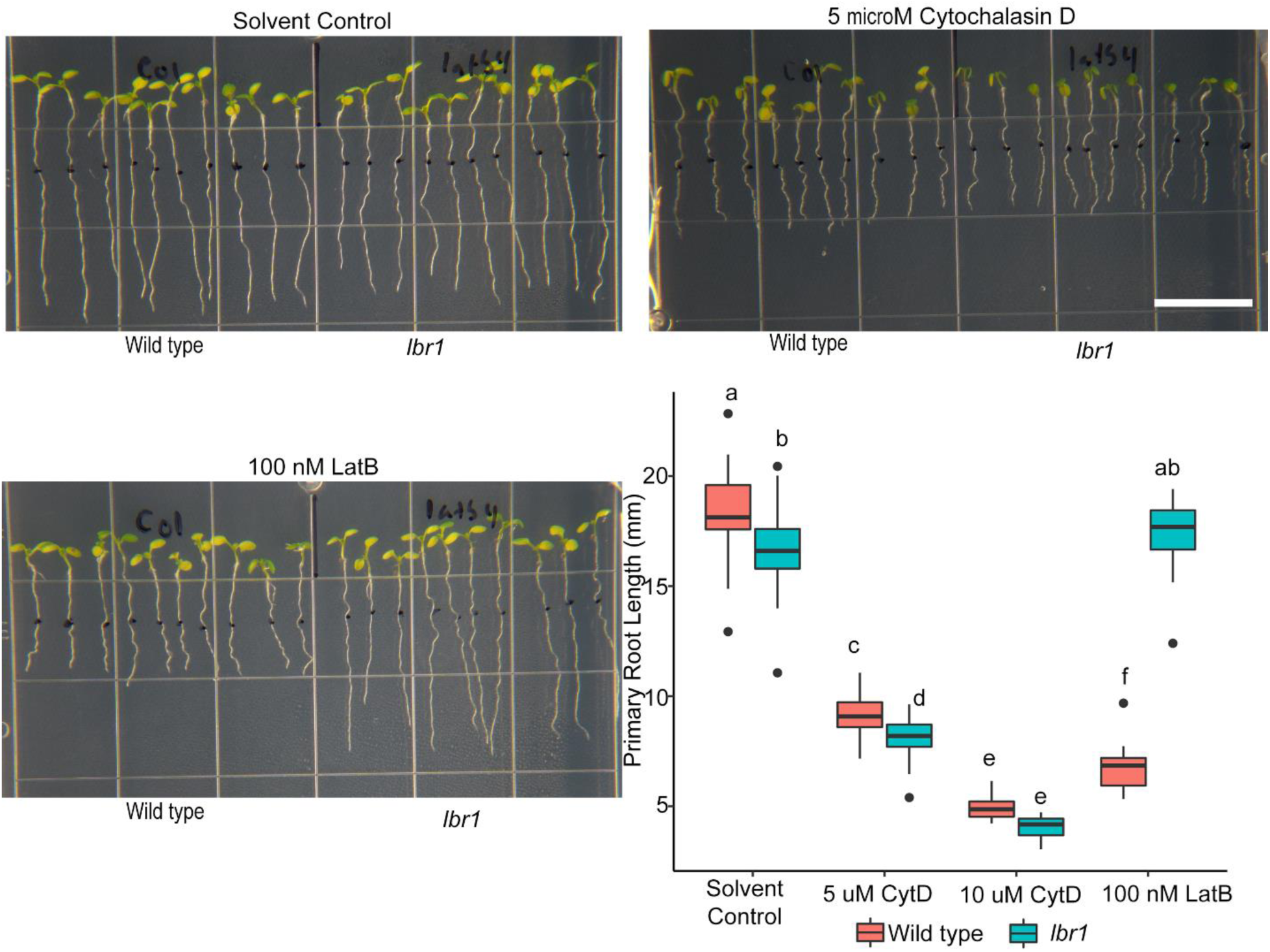
*lbr1* is not resistant to cytochalasin D. Representative images of wild-type and *lbr1* seedlings five days after transplanting to 5 μM cytochalasin D (CytD), 100 nM LatB, or corresponding solvent controls. White bar = 13 mm. Box plots of primary root length of wild-type and *lbr1* seedlings on 0 (solvent control), 5, 10 μM CytD or 100 nM LatB. Box limits indicate 25^th^ and 75^th^ percentiles, horizontal line is the median, and whiskers display minimum and maximum values. Statistical significance was determined by one-way ANOVA. *n* = 30-40 seedlings.

**Fig. S5.**
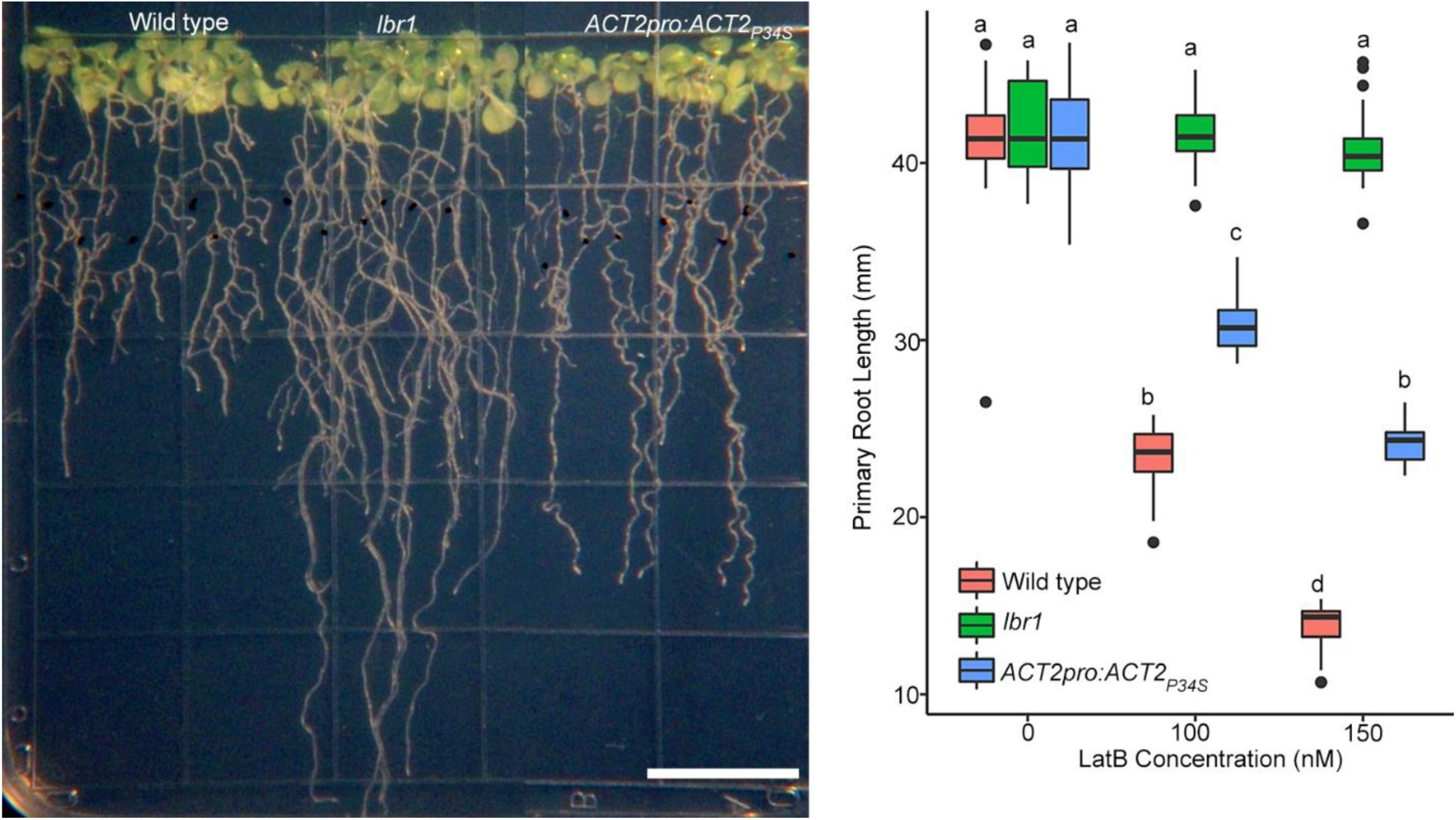
ACT2 P34S confers partial resistance to LatB. Representative images and box plots of wild-type, *lbr1*, and wild type transformed with *ACT2pro:ACT2_P34S_*. Bar = 13 mm. Box limits indicate 25^th^ and 75^th^ percentiles, horizontal line is the median, and whiskers display minimum and maximum values. Statistical significance was determined by one-way ANOVA. *n* = 30-40 seedlings.

## Acknowledgements

BRK acknowledges start-up funds from the Biology Department, University of Scranton.

## Author contributions

JAS, NR, and BRK isolated and characterized the *lbr1* mutant. JAS generated site-directed ACT7 mutants and transgenic lines. JAS, LS, JW, and BRK performed backcrossing, map-based cloning, and next-generation sequencing of *lbr1*. JAS, EBB, and BRK conducted F-actin imaging and analysis. SC, BRK, and EBB generated figures and conducted statistical analyses. SG, EBB, and BRK supervised the research and obtained funding. All authors contributed to the writing of the manuscript.

## Data availability

Seed lines and constructs presented in the paper are available from the corresponding author upon reasonable request.

## References

An YQ, McDowell JM, Huang S, McKinney EC, Chambliss S, Meagher RB. 1996. Strong, constitutive expression of the *Arabidopsis* ACT2/ACT8 actin subclass in vegetative tissues. The Plant Journal 10, 107–121. doi: 10.1046/j.1365-313x.1996.10010107.x

Ayscough KR, Stryker J, Pokala N, Sanders M, Crews P, Drubin DG. 1997. High rates of actin filament turnover in budding yeast and roles for actin in establishment and maintenance of cell polarity revealed using the actin inhibitor latrunculin-A. The Journal of Cell Biology 137, 399–416. doi: 10.1083/jcb.137.2.399

Bibikova TN, Blancaflor EB, Gilroy S. 1999. Microtubules regulate tip growth and orientation in root hairs of *Arabidopsis thaliana*. The Plant Journal 17, 657–665. doi: 10.1046/j.1365-313X.1999.00415.x

Baluška F, Jasik J, Edelmann HG, Salajová T, Volkmann D. 2001a. Latrunculin B-induced plant dwarfism: plant cell elongation is F-actin dependent. Developmental Biology 231, 113–124. doi: 10.1006/dbio.2000.0115

Bascom CS, Hepler PK, Bezanilla M. 2018. Interplay between ions, the cytoskeleton, and cell wall properties during tip growth. Plant Physiology 176, 28–40. doi: 10.1104/pp.17.01466

Baluška F, Busti E, Dolfini S, Gavazzi G, Volkmann D. 2001b. Lilliputian mutant of maize lacks cell elongation and shows defects in organization of actin cytoskeleton. Developmental Biology 236, 478–91. doi: 10.1006/dbio.2001.0333

Blancaflor EB. 2013 Regulation of plant gravity sensing and signaling by the actin cytoskeleton. American Journal of Botany 100, 143–152. doi: 10.3732/ajb.1200283

Baek K, Liu X, Ferron F, Shu S, Korn ED, Dominguez R. 2008. Modulation of actin structure and function by phosphorylation of Tyr-53 and profilin binding. Proceedings of the National Academy of Sciences USA 105, 11748–11753. doi: 10.1073/pnas.0805852105

Chin S, Kwon T, Khan BR, Sparks JA, Mallery EL, Szymanski DB, Blancaflor EB. 2021. Spatial and temporal localization of SPIRRIG and WAVE/SCAR reveal roles for these proteins in actin-mediated root hair development. Plant Cell 33, 2131–2148. doi: 10.1093/plcell/koab115

Chai C, Chin S, Blancaflor EB. 2022. Imaging the cytoskeleton in living plant roots. Methods in Molecular Biology 2364, 139–148. doi: 10.1007/978-1-0716-1661-1_6

Clough SJ, Bent AF. 1998. Floral dip: a simplified method for Agrobacterium-mediated transformation of *Arabidopsis thaliana*. The Plant Journal 16, 735–743. doi: 10.1046/j.1365-313x.1998.00343.x

Dyachok J, Sparks JA, Liao F, Wang YS, Blancaflor EB. 2014. Fluorescent protein-based reporters of the actin cytoskeleton in living plant cells: fluorophore variant, actin binding domain, and promoter considerations. Cytoskeleton (Hoboken) 71, 311–327. doi: 10.1002/cm.21174

Fujita M, Ichinose S, Kiyono T, Tsurumi T, Omori A. 2003. Establishment of latrunculin-A resistance in HeLa cells by expression of R183A D184A mutant β-actin. Oncogene 22, 627– 631. doi: 10.1038/sj.onc.1206173

Gifford ML, Xu G, Dupuy LX, Vissenberg K, Rebetzke G. 2024. Root architecture and rhizosphere-microbe interactions. Journal of Experimental Botany 75, 503–507. doi: 10.1093/jxb/erad488

Gilliland LU, Kandasamy MK, Pawloski LC, Meagher RB. 2002. Both vegetative and reproductive actin isovariants complement the stunted root hair phenotype of the *Arabidopsis act2-1* mutation. Plant Physiology 130, 2199–2209. doi: 10.1104/pp.014068

Gilliland LU, Pawloski LC, Kandasamy MK, Meagher RB. 2003. *Arabidopsis* actin gene ACT7 plays an essential role in germination and root growth. The Plant Journal 33, 319–328. doi: 10.1046/j.1365-313X.2003.01626.x

Gibbon BC, Kovar DR, Staiger CJ. 1999. Latrunculin B has different effects on pollen germination and tube growth. Plant Cell 11, 2349–63. doi: 10.1105/tpc.11.12.2349

Henty-Ridilla JL, Shimono M, Li J, Chang JH, Day B, Staiger CJ. 2013. The plant actin cytoskeleton responds to signals from microbe-associated molecular patterns. PLoS Pathogens 9, e1003290. doi: 10.1371/journal.ppat.1003290

Hiles R, Rogers A, Jaiswal N, Zhang W, Butchacas J, Merfa MV, Klass T, Barua P, Thirumalaikumar VP, Jacobs JM, Staiger CJ, Helm M, Iyer-Pascuzzi AS. 2024. A *Ralstonia solanacearum* type III effector alters the actin and microtubule cytoskeleton to promote bacterial virulence in plants. PLoS Pathogens 20, e1012814. doi: 10.1371/journal.ppat.1012814

Hou G, Mohamalawari DR, Blancaflor EB. 2003. Enhanced gravitropism of roots with a disrupted cap actin cytoskeleton. Plant Physiology 131, 1360–1373. doi: 10.1104/pp.014423

Hou G, Kramer VL, Wang Y-S, Chen R, Perbal G, Gilroy S, Blancaflor EB. 2004. The promotion of gravitropism in *Arabidopsis* roots upon actin disruption is coupled with the extended alkalinization of the columella cytoplasm and a persistent lateral auxin gradient. The Plant Journal 39, 113–125. doi: 10.1111/j.1365-313X.2004.02114.x

Iwabuchi K, Ohnishi H, Tamura K, Fukao Y, Furuya T, Hattori K, Tsukaya H, Hara-Nishimura I. 2019. ANGUSTIFOLIA Regulates Actin Filament Alignment for Nuclear Positioning in Leaves. Plant Physiology 179, 233–247. doi: 10.1104/pp.18.01150

Kandasamy MK, McKinney EC, Meagher RB. 2010. Differential sublocalization of actin variants within the nucleus. Cytoskeleton (Hoboken) 67, 729–743. doi: 10.1002/cm.20484

Kandasamy MK, McKinney EC, Meagher RB. 2009. A single vegetative actin isovariant overexpressed under the control of multiple regulatory sequences is sufficient for normal *Arabidopsis* development. Plant Cell 21, 701–718. doi: 10.1105/tpc.108.061960

Kandasamy MK, Gilliland LU, McKinney EC, Meagher RB. 2001. One plant actin isovariant, ACT7, is induced by auxin and required for normal callus formation. Plant Cell 13, 1541–1554. doi: 10.1105/tpc.010026

Koboldt DC, Zhang Q, Larson DE, Shen D, McLellan MD, Lin L, Miller CA, Mardis ER, Ding L, Wilson RK. 2012. VarScan 2: somatic mutation and copy number alteration discovery in cancer by exome sequencing. Genome Research 22, 568–576. doi: 10.1101/gr.129684.111

Kudrimoti S, Ahmed SA, Daga PR, Wahba AE, Khalifa SI, Doerksen RJ, Hamann MT. 2009. Semisynthetic latrunculin B analogs: studies of actin docking support a proposed mechanism for latrunculin bioactivity. Bioorganic & Medicinal Chemistry 17, 7517–7522. doi: 10.1016/j.bmc.2009.09.012

Kato T, Morita MT, Tasaka M. 2010. Defects in dynamics and functions of actin filament in *Arabidopsis* caused by the dominant-negative actin fiz1-induced fragmentation of actin filament. Plant & Cell Physiology 51, 333–338. doi: 10.1093/pcp/pcp189

Leontovyčová H, Kalachova T, Trdá L, Pospíchalová R, Lamparová L, Dobrev PI, Malínská K, Burketová L, Valentová O, Janda M. 2019. Actin depolymerization is able to increase plant resistance against pathogens via activation of salicylic acid signalling pathway. Scientific Reports 9, 1–10. doi: 10.1038/s41598-019-46465-5

Li H, Durbin R. 2009. Fast and accurate short read alignment with Burrows-Wheeler transform. Bioinformatics, vol 25. pg. 1754–1760

Lenth RV. 2016. Least-squares means: The R Package lsmeans. Journal of Statistical Software 69, 1. doi: 10.18637/jss.v069.i01

Leontovyčová H, Kalachova T, Janda M. 2020. Disrupted actin: a novel player in pathogen attack sensing? New Phytologist 227,1605–1609. doi: 10.1111/nph.16584

Mao H, Nakamura M, Viotti C, Grebe M. 2016. A framework for lateral membrane trafficking and polar tethering of the PEN3 ATP-binding cassette transporter. Plant Physiology 172, 2245– 2260. doi: 10.1104/pp.16.01252

Maeda K, Sasabe M, Hanamata S, Machida Y, Hasezawa S, Higaki T. 2020. Actin filament disruption alters phragmoplast microtubule dynamics during the initial phase of plant cytokinesis. Plant Cell Physiology 61, 445–456. doi: 10.1093/pcp/pcaa003

McLean BG, Huang S, McKinney EC, Meagher RB. 1990. Plants contain highly divergent actin isovariants. Cell Motil Cytoskeleton 17, 276–290. doi: 10.1002/cm.970170403

Meagher RB. 1991. Divergence and differential expression of actin gene families in higher plants. International Review of Cytology 125, 139–163. doi: 10.1016/s0074-7696(08)61218-8

McDowell JM, Huang S, McKinney EC, An YQ, Meagher RB. 1996a. Structure and evolution of the actin gene family in *Arabidopsis thaliana*. Genetics 142, 587–602. doi: 10.1093/genetics/142.2.587

Meagher RB, McKinney EC, Vitale AV. 1999. The evolution of new structures: clues from plant cytoskeletal genes. Trends In Genetics 15, 278–283. doi: 10.1016/s0168-9525(99)01759-x

McDowell JM, An YQ, Huang S, McKinney EC, Meagher RB. 1996b. The *Arabidopsis* ACT7 actin gene is expressed in rapidly developing tissues and responds to several external stimuli. Plant Physiology 111, 699–711. doi: 10.1104/pp.111.3.699

Meagher RB, McKinney EC, Kandasamy MK. 2000. The significance of diversity in the plant actin gene family: studies in Arabidopsis. In: Staiger CJ, Baluška F, Volkmann D, Barlow P, eds, Actin: A Dynamic Framework for Multiple Plant Cell Functions. Dordrecht, The Netherlands: Kluwer Academic Publishers, pp. 3–27.

Morton WM, Ayscough KR, McLaughlin PJ. 2000. Latrunculin alters the actin-monomer subunit interface to prevent polymerization. Nature Cell Biology 2, 376–378. doi: 10.1038/35014075

Mathur J, Spielhofer P, Kost B, Chua N-H. 1999. The actin cytoskeleton is required to elaborate and maintain spatial patterning during trichome cell morphogenesis in *Arabidopsis thaliana*. Development 126, 5559–5568. doi: 10.1242/dev.126.24.5559

Nishimura T, Yokota E, Wada T, Shimmen T, Okada K. 2003. An *Arabidopsis* ACT2 dominant-negative mutation, which disturbs F-actin polymerization, reveals its distinctive function in root development. Plant Cell Physiology 44, 1131–1140. doi: 10.1093/pcp/pcg158

Numata T, Sugita K, Ahamed Rahman A, Rahman A. 2022. Actin isovariant ACT7 controls root meristem development in *Arabidopsis* through modulating auxin and ethylene responses. Journal of Experimental Botany 73, 6255–6271. doi: 10.1093/jxb/erac280

Paez-Garcia A, Sparks JA, de Bang L, Blancaflor EB. 2018. Plant Actin Cytoskeleton: New Functions from Old Scaffold. In: Sahi V., Baluška F. (eds) Concepts in Cell Biology - History and Evolution. Plant Cell Monographs, vol 23. pp. 103–137, Springer, Cham doi: 10.1007/978-3-319-9944-8_6

Pospich S, Merino F, Raunser S. 2020. Structural effects and functional implications of phalloidin and jasplakinolide binding to actin filaments. Structure 28, 437–449. doi: 10.1016/j.str.2020.01.014

Ringli C, Baumberger N, Diet A, Frey B, Keller B. 2002. ACTIN2 is essential for bulge site selection and tip growth during root hair development of *Arabidopsis*. Plant Physiology 129, 464–472. doi: 10.1104/pp.005777

Rosero A, Žárský V, Cvrčková F. 2013. AtFH1 formin mutation affects actin filament and microtubule dynamics in *Arabidopsis thaliana*. Journal of Experimental Botany 64, 585–597. doi: 10.1093/jxb/erv473

Reikofski J, Tao BY. 1992. Polymerase chain reaction (PCR) techniques for site-directed mutagenesis. Biotechnology Advances 10, 535–547. doi: 10.1016/0734-9750(92)91451-J

R Core Team. 2019. R: a language and environment for statistical computing. In R Foundation for Statistical Computing. Vienna, Austria

Staiger CJ, Sheahan MB, Khurana P, Wang X, McCurdy DW, Blanchoin L. 2009. Actin filament dynamics are dominated by rapid growth and severing activity in the *Arabidopsis* cortical array. The Journal of Cell Biology 184, 269–280. doi: 10.1083/jcb.200806185

Sparks JA, Kwon T, Renna L, Liao F, Brandizzi F, Blancaflor EB. 2016. HLB1 is a tetratricopeptide repeat domain-containing protein that operates at the intersection of the exocytic and endocytic pathways at the TGN/EE in *Arabidopsis*. Plant Cell 28, 746–769. doi: 10.1105/tpc.15.00794

Sun L, Ge Y, Sparks JA, Robinson ZT, Cheng X, Wen J, Blancaflor EB. 2019. TDNAscan: A software to identify complete and truncated T-DNA insertions. Frontiers in Genetics 10, 685. doi: 10.3389/fgene.2019.00685

Seethepalli A, Dhakal K, Griffiths M, Guo H, Freschet GT, York LM. 2021. RhizoVision Explorer: open-source software for root image analysis and measurement standardization. AoB PLANTS 13, plab056. doi: 10.1093/aobpla/plab056

Seethepalli A, York LM. 2020. RhizoVision Explorer - Interactive software for generalized root image analysis designed for everyone (Version 2.0.3). Zenodo. doi: 10.5281/zenodo.4095629

Schliwa, M. 1982. Action of cytochalasin D on cytoskeletal networks. The Journal of Cell Biology, 92, 79–91. doi: 10.1083/jcb.92.1.79

Spector I, Shochet NR, Kashman Y, Groweiss A.1983. Latrunculins: novel marine toxins that disrupt microfilament organization in cultured cells. Science 219, 493–495. doi: 10.1126/science.6681676

Skolnick J, Gao M, Zhou H, Singh S. 2021. AlphaFold 2: Why It Works and Its Implications for Understanding the Relationships of Protein Sequence, Structure, and Function. Journal of chemical information and modeling 61, 4827–4831. doi: 10.1021/acs.jcim.1c01114

Tsang I, Atkinson JA, Rawsthorne S, Cockram J, Leigh F. 2024. Root hairs: an underexplored target for sustainable cereal crop production. Journal of Experimental Botany 75, 5484–5500. doi: 10.1093/jxb/erae275

Takatsuka H, Higaki T, Umeda M. 2018. Actin reorganization triggers rapid cell elongation in roots. Plant Physiology 178, 1130–1141. doi: 10.1104/pp.18.00557

Vaškebová L, Šamaj J, Ovecka M. 2018. Single-point *ACT2* gene mutation in the *Arabidopsis* root hair mutant *der1-3* affects overall actin organization, root growth and plant development. Annals of Botany 122, 889–901. doi: 10.1093/aob/mcx180

Wang K, Li M, Hakonarson H. 2010. ANNOVAR: functional annotation of genetic variants from high-throughput sequencing data. Nucleic Acids Research 38, e164. doi: 10.1093/nar/gkq603

Yang H, Wang K. 2015. Genomic variant annotation and prioritization with ANNOVAR and wANNOVAR. Nature Protocols 10, 1556–1566. doi: 10.1038/nprot.2015.105

Wang YS, Yoo CM, Blancaflor EB. 2008. Improved imaging of actin filaments in transgenic *Arabidopsis* plants expressing a green fluorescent protein fusion to the C- and N-termini of the fimbrin actin-binding domain 2. The New Phytologist 177, 525–536. doi: 10.1111/j.1469-8137.2007.02261.x

Wickham H. 2016. ggplot2: Elegant Graphics for Data Analysis. Springer-Verlag New York.

Yoo C-M, Quan L, Cannon AE, Wen J, Blancaflor EB. 2012. AGD1, a class 1 ARF-GAP, acts in common signaling pathways with phosphoinositide metabolism and the actin cytoskeleton in controlling *Arabidopsis* root hair polarity. The Plant Journal 69, 1064–1076. doi: 10.1111/j.1365-313X.2011.04856.x

Zhu J, Bailly A, Zwiewka M, Sovero V, Di Donato M, Ge P, Oehri J, Aryal B, Hao P, Linnert M, et al. 2016. TWISTED DWARF1 mediates the action of auxin transport inhibitors on actin cytoskeleton dynamics. Plant Cell 28, 930–948. doi: 10.1105/tpc.15.00726

